# A functional analysis of action potential generation in the axon initial segment in two autism mouse models with loss-of-function mutations of the Na_v_1.2 sodium channel

**DOI:** 10.64898/2026.11.24.768870

**Authors:** Fatima Abbas, Elisabetta Giacalone, Laila Ananda Blömer, Ömer Yusuf İpek, Domenico Tegolo, Massimo Mantegazza, Michele Migliore, Marco Canepari

**Author notes:** Address of the submitting and corresponding author: Marco Canepari, Laboratoire Interdisciplinaire de Physique (UMR 5588), Bat. E45, 140 avenue de la physique, Domaine univ., 38402 St Martin d’Hères cedex, France.

## Abstract

Loss-of-function (LoF) mutations of the *SCN2A* gene, coding for Na_v_1.2 Na^+^ channels, are among the leading genetic contributors to autism spectrum disorders and intellectual disability (ASD/ID). The haploinsufficient *Scn2a* knock-out mouse (*Scn2a^+/−^*), displaying ASD/ID-like phenotypes, is commonly used as model for this channelopathy, although this specific mutation has not been identified in patients. We investigated the action potential (AP) generation in the axon initial segment (AIS) of layer-5 neocortical pyramidal neurons in wild-type (*Scn2a^+/+^*) mice, in *Scn2a^+/−^* mice and in mice carrying the missense mutation L1314P (*Scn2a^+/L1314P^*), identified in an ASD/ID patient. We recorded somatic action potentials (APs), and imaged Na^+^, Ca^2+^ and APs in the AIS, in females during postnatal (P) P21-40. During maturation from P21-24 to P30-40: (1) the ratio between Na^+^ influx in proximal and distal AIS sites (*R_p/d_*) increased; (2) the “*delay”* between distal and proximal Ca^2+^ currents increased; (3) the half-width of the AP (*AP_wid_*) in the soma decreased. We revealed that this maturation process is different in both *Scn2a^+/−^* and *Scn2a^+/L1314P^* mice, but in divergent manner. With respect to *Scn2a^+/+^*, *Scn2a^+/-^* cells were characterised by smaller *R_p/d_*and *delay* values at P30-40, and wider somatic *AP_wid_* at P21-24, whereas *Scn2a^+/L1314P^* cells by shorter AIS *AP_wid_*at P21-24. Using a biophysical model, mimicking maturation changes in terms of Na_v_1.2, of a K^+^ channel and of voltage-gated Ca^2+^ channels, we reproduced the ensemble of experimental results, establishing that *Scn2a^+/-^* cells exhibit a combination of Na_v_1.2 LoF and delayed development, whereas *Scn2a^+/L1314P^* cells are characterised by AIS disruption involving strongly increased functional K^+^ channels. We conclude that diverse Na_v_1.2 LoF cases lead to different developmental paths involving other channels.

## Introduction

The gene *Scn2a* encodes the voltage-gated Na^+^ channel (VGNC) Na_v_1.2 α-subunit, which is one of the principal VGNC isoforms in the brain [1]. In many myelinated neurons, Na_v_1.2 is involved in action potential (AP) generation in the axon initial segment (AIS) [2,3,4], in axonal propagation in developmental Ranvier nodes [5] and in dendritic back-propagation. Mutations of *Scn2a* have been identified as prominent causes for a large spectrum of neurodevelopmental disorders [6]. *Scn2a*/Na_v_1.2 mutations can be broadly classified into variants that induce *gain-of-function* (GoF) and variants that induce *loss of function* (LoF) on the resulting channel [1,7,8]. GoF mutations cause neonatal-early infantile developmental and epileptic encephalopathy (NEIDEE) or milder self-limited (familial) neonatal-infantile epilepsy (SeLFNIE), whereas LoF mutations cause late infantile-childhood developmental and epileptic encephalopathy (ICDEE), which are among the leading genetic contributors to autism spectrum disorder and intellectual disability (ASD/ID) [9,10,11]. Na_v_1.2 is expressed in many cellular types and, in the cortex, prominently in pyramidal neurons [12,13]. Thus, GoF mutations are thought to increase neuronal excitability, consistent with their association with epilepsy, whereas LoF mutations are expected to reduce excitability, but the mechanisms linking Na_V_1.2 LoF to ICDEE or ASD/ID remain incompletely understood. In line with this framework, haploinsufficient heterozygous *Scn2a* knock-out mice (*Scn2a^+/−^*) display ASD/ID-like phenotypes [14,15], with reduced excitability and synaptic transmission in cortical [16] and hippocampal [17] pyramidal neurons. Notably, some LoF missense mutations can induce other effects than reduced current. For instance, *Scn2a*^+/K1422E^ knock-in mice, carrying a missense variant associated to ICDEE with developmental delay, infantile spasms and ASD, exhibits Na_v_1.2 currents with larger permeability to Ca^2+^ and resistance to tetrodotoxin, as well as abnormal signalling in AP generation in the AIS [18]. More generally, the pathogenic consequences of a variant may arise not only from its direct effects on channel function but also from secondary responses. Consistently, severe Na_v_1.2 deficiency in mice has been shown to paradoxically increase neuronal excitability, attributed to the interplay between Na^+^ and K^+^ channels or down-regulation of voltage-gated K^+^ channels (VGKCs) [19,20]. However, secondary responses may also involve broader cellular, developmental, or circuit-level dysfunctions that cannot be fully explained by the properties of the affected channel alone. A feature that can be altered is AIS targeting of VGNCs [21,22], causing functional remodeling of this important cellular sub-compartment, which is governed by scaffold proteins such as ankyrin-G (AnkG) [23]. Evidence is accumulating for AIS dysfunction in several pathological conditions [24]. The AIS is often explored using structural life imaging techniques [25], while functional approaches are necessary to understand the impact of remodeling in cases of genetic alterations. As an example, Na^+^ and Ca^2+^ imaging in the AIS, in combination with somatic patch clamp recording, was used to investigate selective suppression of Na_v_1.6 in neocortical layer-5 (L5) pyramidal neurons [26]. Since AP generation is a multistep process evolving on a time window of ∼10-1000 µs, functional imaging should be ideally performed at this time resolution. We recently developed an imaging technique to resolve the kinetics of Na^+^ transients associated with an AP in the AIS [27], and we combined this technique with Ca^2+^ and membrane potential (V_m_) imaging at the same temporal resolution to investigate Na_v_1.2 function in L5 pyramidal neurons [28]. Here, we report a comparative study of haploinsufficient *Scn2a*^+/−^ knock-out mice and *Scn2a*^+/L1314P^ mice carrying the L1314P missense variant identified in a patient with severe ASD/ID [29], using the same experimental approach applied in our previous work. L1314P has been shown to impair plasma membrane trafficking of Na_V_1.2 and, based on *in vitro* evidence, to exert dominant-negative effects that further reduce the membrane expression of co-expressed wild-type Na_V_1.2 channels [30]. Both mouse lines display specific ASD/ID-like phenotypes during the postnatal period spanning P20 to P90. Thus, our analysis focussed on a period within this age window. To achieve robust statistical power, we performed imaging experiments from 350 cells from 156 female mice in the time window P21-P40, and additionally patch-clamp recordings of somatic AP from 416 cells in 41 female mice. We initially focussed on optical signals directly linked to Na_v_1.2 function, based on the hypothesis that both mutant strains would exhibit reduced functional channel in the plasma membrane. However, our attention progressively shifted as it became apparent that the observed differences in AP waveform could not be accounted for solely by Na_v_1.2 dysfunction. We found that both mutated strains differ from the wild-type (*Scn2a^+/+^*) mouse, but in divergent manner. To understand the mechanisms underlying this divergence, we developed a minimal biophysical model that captures the key phenotypic differences between the two strains.

## Results

### Na^+^ influx and Ca^2+^ current in the AIS of *Scn2a^+/-^* and *Scn2a^+/L1314P^* mice

The analysis of signals in *Scn2a^+/-^* and *Scn2a^+/L1314P^*mice began with the measurement of Na^+^ influx associated with the generation of an AP in the AIS of L5 pyramidal neurons of the somato-sensory cortex, using a quantitative method that we developed in the laboratory for wild-type mice [27]. Here, we performed new experiments in control *Scn2a^+/+^* female mice, from the same litters of transgenic animals, associating with each cell the postnatal day in the range P21-P40. In the two representative examples shown in Figure 1A, an AP evoked in the soma and the associated Na^+^ concentration changes from three sites of the AIS are reported from a cell at P21 (left) and from another cell at P30 (right). The distal (*d*, green traces), medial (*m*, red traces) and proximal (*p*, blue traces) sites correspond to 5µm-long regions in the AIS at 30-40 µm, 15-25 µm and 0-10 µm distances from the soma. In the P21 cell, Na^+^ transients were comparable in size at *d* and *m* sites whereas the signal at the *p* site was smaller. In the P30 cell, in contrast, Na^+^ transients were comparable in size at *d* and *p* sites whereas the signal at the *m* site was larger. To quantify the size of the signal, we fitted the raw traces with a model function. We then extrapolated the Na^+^ current (*I_Na_*) by calculating the time derivative and by estimating the AIS surface at each site [27]. In these two examples, the smallest *I_Na_*was at *p* in the P21 cell, and at *d* in the P30 cell respectively. The quantification of the *I_Na_*, however, depends on the undersampled measurement of the fast transient and on the imprecise estimate of the AIS morphology. Thus, we restricted our analysis to the size of the transient fits, which is not affected by these errors, and we focussed on *d* and *p* sites. We previously showed that the partially-selective Na_v_1.2 blocker G1G4-huwentoxin-IV inhibits Na^+^ influx more strongly in the proximal part of the AIS, with respect to the distal part [28] where Na_v_1.6 is preferentially expressed [12]. Thus, whereas the absolute amplitude of the Na^+^ concentration change is variable from cell to cell, the ratio between the signals in *p* and *d* (*R_p/d_*) is an indication of the Na_v_1.2 signal with respect to the Na_v_1.6 signal. In the two examples in Figure 1A, the values of *R_p/d_* were 0.63 at P21 and 1.02 at P30. We measured Na^+^ influx in 29 cells from 13 *Scn2a^+/+^* females and the values of *R_p/d_* against the age are reported in Supplemental Figure 1. We observed that *R_p/d_* has tendency of increasing with age with values always <1 in the 13 cells belonging to the youngest group (P21-24) and either positive or negative values closer to 1 in the oldest group (P30-40), consistently with what we previously reported [27,28]. Since this result suggests a refinement of Na_v_1.2 expression in the examined maturation period, in this study we linked signals to three age groups of females: the entire age range group (P21-40); the young group (P21-24); and the old group (P30-40). The two examples shown in Figure 1B report the results of the same type of experiment performed in a cell at P21 (left) and in another cell at P30 (right) from *Scn2a^+/-^* female mice. In both cells, Na^+^ signals were smaller in *p* with respect to *d* and values of *R_p/d_* were 0.37 and 0.65 at P21 and P30 respectively. Values of *R_p/d_* against the age in 31 cells from 12 *Scn2a^+/-^* mice, reported again in Supplemental Figure 1, were always <1. Finally, the two examples shown in Figure 1C report the results in a cell at P21 (left) and in another cell at P35 (right) from *Scn2a^+/L1314P^*females.

**Figure 1.**
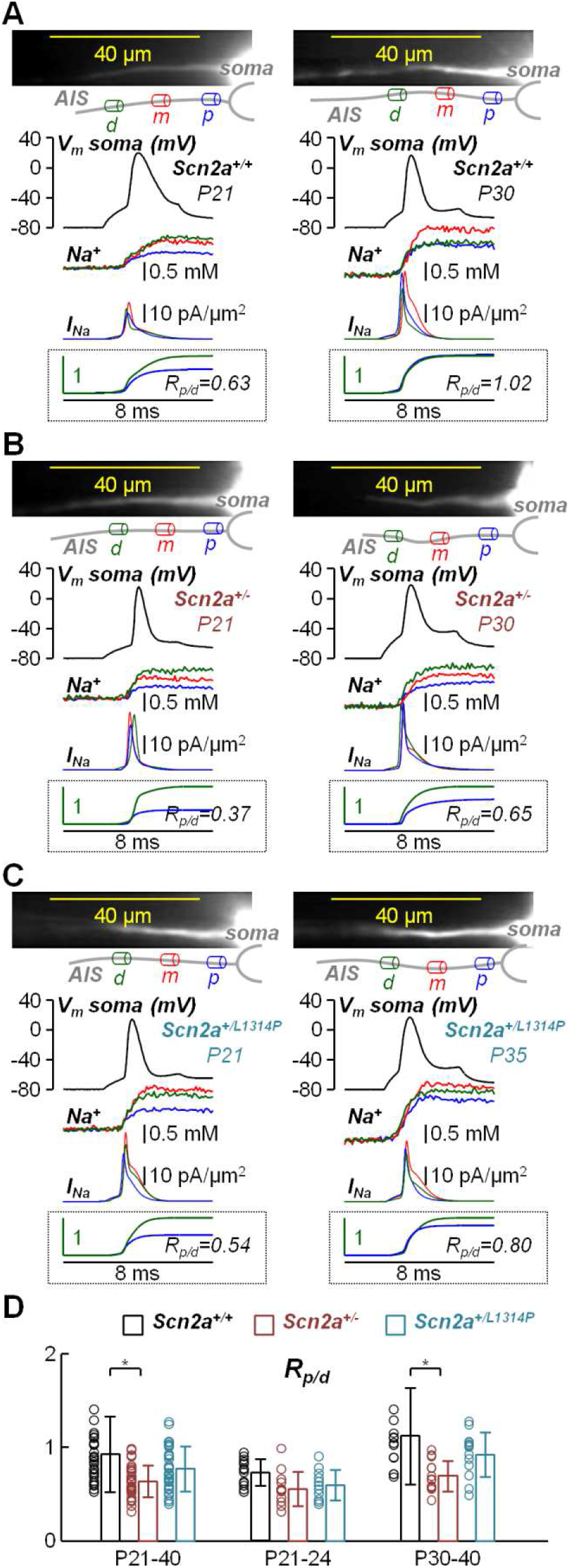
Na^+^ influx in the AIS of *Scn2a^+/+^, Scn2a^+/-^* and *Scn2a^+/L1314P^* L5 pyramidal neurons. (A) On the top-left, AIS of a *Scn2a^+/+^* L5 pyramidal neuron at P21 in the recording position with reconstruction indicating a distal (*d*, green cylinder), a medial (*m*, red cylinder) and a proximal (*p*, blue cylinder) site. On the middle-left, an AP with associated Na^+^ transients from the three sites illustrated with the same colour of the corresponding cylinders above. Below, Na^+^ currents (*I_Na_*) derived from the fits of the Na^+^ transients. Inside the dotted inset, fits of the Na^+^ transients in the *d* and *p* sites normalised to the maximum of the *d* fit; the value of the ratio between the *p* and the *d* maxima (*R_p/d_*) is reported. On the right, same measurements and analysis for another *Scn2a^+/+^* cell at P30. Notice the difference between the *R_p/d_* values in the two neurons. (B) Same measurements and analysis for two *Scn2a^+/-^* L5 pyramidal neurons, one at P21 (left) and the other at P30 (right). Notice that *R_p/d_*is less than one in both cells. (C) Same measurements and analysis for two *Scn2a^+/L1314P^* L5 pyramidal neurons, one at P21 (left) and the other at P35 (right). (D) Scatter plots and mean ± SD of *R_p/d_* values for the three strains and the three age groups. In *Scn2a^+/+^* cells, *R_p/d_* was 0.92±0.40 at P21-40 (N=29), 0.73±0.14 at P21-24 (N=13) and 1.12±0.51 at P30-40 (N=13). In *Scn2a^+/-^* cells, *R_p/d_* was 0.64±0.17 at P21-40 (N=31), 0.56±0.18 at P21-24 (N=12) and 0.69±0.16 at P30-40 (N=13). In *Scn2a^+/L1314P^* cells, *R_p/d_* was 0.77±0.24 at P21-40 (N=29), 0.59±0.16 at P21-24 (N=11) and 0.92±0.24 at P30-40 (N=13). “*” indicates that the result of the two-sample ks-test between two groups gave P<0.01 and SP>0.8.

*R_p/d_* was 0.54 in the cell at P21 and 0.80 in the cell at P35. The values of *R_p/d_* against the age in 29 cells from 12 *Scn2a^+/L1314P^* females, also reported in Supplemental Figure 1, were qualitatively more similar to those of *Scn2a^+/+^* cells, with values >1 measured in 6 cases in the older group. To assess potential differences among the different groups, we used the two-sample Kolmogorov-Smirnov (ks) test and we established that two groups are different when both the P value is <0.01 and the Statistical Power (SP) is >0.8 (see Methods). Figure 1D shows the scatter plots of *R_p/d_*values for the three strains and the three age groups, together with the mean ± SD. *R_p/d_* values in *Scn2a^+/-^* cells were smaller than those in *Scn2a^+/+^* cells both at P21-40 (P=0.0023 and SP=0.952) and at P30-40 (P=0.0076 and SP=0.821). Interestingly, in the old group, the ks-test performed between *Scn2a^+/-^* and *Scn2a^+/L1314P^* cells gave P=0.028 and SP=0.814. All values of the statistical analysis of Na^+^ influx experiments are reported in Supplemental Table 1. Altogether, the results of the analysis suggest a clear loss of Na^+^ influx in *Scn2a^+/-^* cells, but not in *Scn2a^+/L1314P^* cells.

Na_v_1.2 is also permeable to Ca^2+^ [31] and we recently demonstrated that Ca^2+^ influx *via* Na_v_1.2 can be distinguished from that *via* voltage-activated Ca^2+^ channels (VGCCs) by the different kinetics [28]. In the two representative examples shown in Figure 2A, fractional changes of Ca^2+^ fluorescence (ΔF/F_0_), associated with an AP evoked in the soma, are reported from *d* and *p* sites in two *Scn2a^+/+^* cells at P22 (left) and at P31 (right). The kinetics of the ΔF/F_0_ signal is composed of two components of Ca^2+^ influx, mediated by Na_v_1.2 and VGCCs respectively. We have previously shown that the VGCC component is more prominent in the proximal AIS [28]. Thus, after deriving the kinetics of the total Ca^2+^ current (*I_Ca_*) using a previously described approach [32,33], the peak of the *I_Ca_* signal at proximal sites is consistently delayed with respect to that at distal sites [28]. The variable “*delay*”, providing information on Na_v_1.2 Ca^2+^ influx with respect to VGCC Ca^2+^ influx, was 200 µs (2 frames) and 500 µs (5 frames) in the P22 cell and in the P31 cell respectively. We measured Ca^2+^ influx in 38 cells from 16 *Scn2a^+/+^* females and the histograms of *delay*, for the three groups P21-40, P21-24 and P30-40, are reported in Supplemental Figure 2. Since *delay* is a discrete variable, results were assessed by applying the more appropriate two-sample Wilcoxon-Mann-Whitney (wmw) test, establishing that two groups are different when both P <0.01 and Standardised Effect Size (SES) >0.5. The *delay* in the young group was smaller than that in the old group (P=0.0012 and SES=0.66) and the statistics of *delay* at P30-40 (383 ± 111 µs) was consistent with that previously reported [28]. The two examples shown in Figure 2B report the results of Ca^2+^ recordings performed in a cell at P23 (left) and in another cell at P37 (right) from *Scn2a^+/-^* females. The histograms of *delay* from 39 cells from 15 *Scn2a^+/-^* females for the three age groups are also reported in Supplemental Figure 2. In these mice, a difference in *delay* between the young group and the old group was not detected (P=0.1709 and SES=0.23). Finally, the two examples shown in Figure 2C report the results of Ca^2+^ recordings performed in a cell at P23 (left) and in another cell at P30 (right) from two *Scn2a^+/L1314P^* females. The histograms of *delay* from 37 cells from 10 *Scn2a^+/L1314P^*females are reported in Supplemental Figure 2 and, as in the case of *Scn2a^+/-^* mice, a difference in *delay* between the young group and the old group was not observed (P=0.1268 and SES=0.29). Figure 2D shows mean ± SD of *delay* for the three strains and the three age groups. The *delay* in *Scn2a^+/-^* cells was smaller than that in *Scn2a^+/+^* cells both at P21-40 (P=1.25·10^-5^ and SES=0.55) and at P30-40 (P=1.09·10^-5^ and SES=0.75), and in the old group it was also smaller than that in *Scn2a^+/L1314P^* cells (P=0.0018 and SES=0.51). All values of the statistical analysis of Ca^2+^ influx experiments are reported in Supplemental Table 2. The results from the analysis of Na^+^ influx and of Ca^2+^ currents are apparently consistent with an effective decrease of functional Na_v_1.2 channels in *Scn2a^+/-^* cells with respect to the wild-type strain. A different scenario, however, seems to occur in *Scn2a^+/L1314P^* cells.

**Figure 2.**
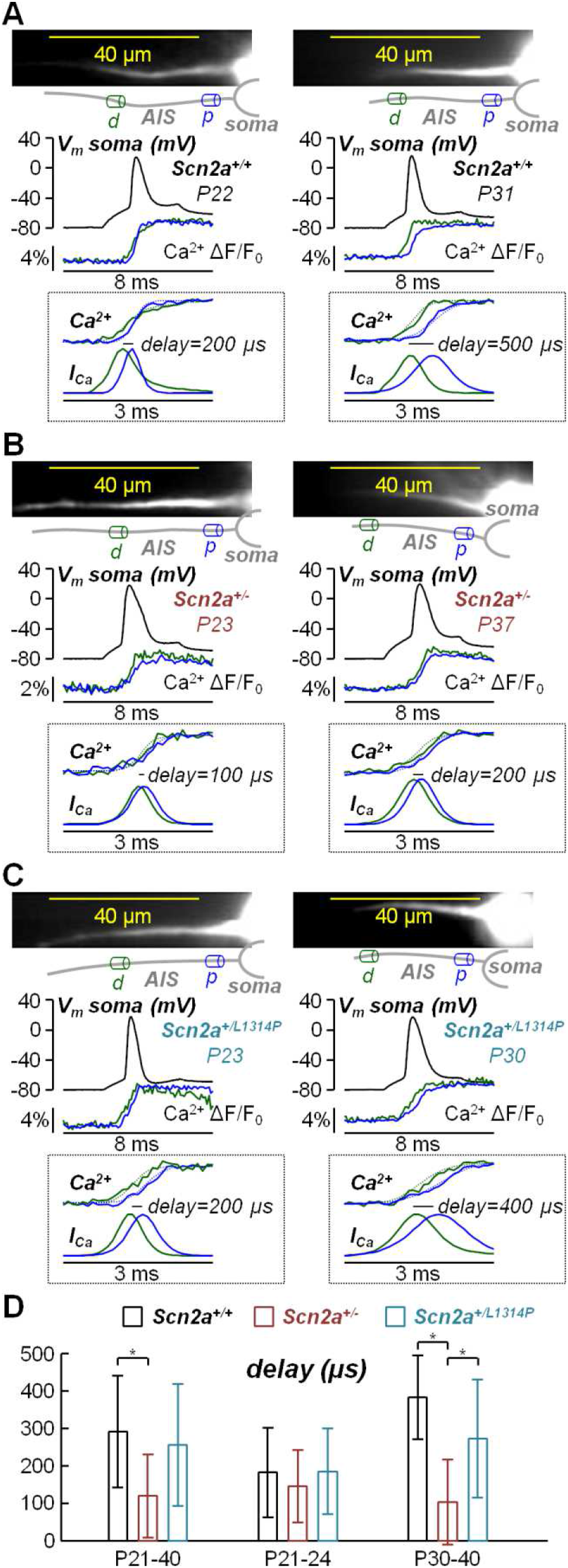
Ca^2+^ influx in the AIS of *Scn2a^+/+^, Scn2a^+/-^* and *Scn2a^+/L1314P^* L5 pyramidal neurons. (A) On the top-left, AIS of a *Scn2a^+/+^* L5 pyramidal neuron at P22 in the recording position with reconstruction indicating a distal (*d*, green cylinder) and a proximal (*p*, blue cylinder) site. On the middle-left, an AP with associated Ca^2+^ transients from the two sites illustrated with the same colour of the corresponding cylinders above. Inside the dotted inset, the same Ca^2+^ transients over 3 ms with fits (dotted lines) and, below, corresponding derived Ca^2+^ currents (*I_Ca_*); *delay* between *I_Ca_* peak at d and at p (200 µs) is indicated. On the right, same measurements and analysis for another *Scn2a^+/+^* cell at P31. In this case *delay* was 500 µs. (B) Same measurements and analysis for two *Scn2a^+/-^* L5 pyramidal neuron, one at P23 (left) and the other at P37 (right). The values of *delay* were 100 µs and 200 µs in the younger and older cells respectively. (C) Same measurements and analysis for two *Scn2a^+/L1314P^* L5 pyramidal neurons, one at P23 (left) and the other at P30 (right). The values of *delay* were 200 µs and 400 µs in the younger and older cells respectively. (D) Mean ± SD of *delay* for the three strains and the three age groups. In *Scn2a^+/+^* cells, *delay* was 292±150 µs at P21-40 (N=38), 183±119 µs at P21-24 (N=12) and 383±111 µs at P30-40 (N=12). In *Scn2a^+/-^* cells, *delay* was 121±110 µs at P21-40 (N=39), 146±97 µs at P21-24 (N=13) and 105±113 µs at P30-40 (N=22). In *Scn2a^+/L1314P^* cells, *delay* was 257±163 µs at P21-40 (N=37), 185±114 µs at P21-24 (N=13) and 273±160 µs at P30-40 (N=15). “*” indicates that the result of the two-sample wmw-test between two groups gave P<0.01 and SES>0.5 and.

### Somatic APs in *Scn2a^+/-^* and *Scn2a^+/L1314P^* mice

Next, we measured somatic APs in *Scn2a^+/-^* and *Scn2a^+/L1314P^* cells and compared the results with those obtained in control *Scn2a^+/+^* cells. We performed experiments in brain slices from 15 *Scn2a^+/+^* female mice, 12 *Scn2a^+/-^* female mice and 14 *Scn2a^+/L1314P^* female mice, at different postnatal days, and we obtained data from 142 *Scn2a^+/+^* cells, 138 *Scn2a^+/-^* cells and 136 *Scn2a^+/L1314P^* cells (Supplemental Table 3). In these experiments, we elicited APs by somatic injection of a current pulse of 50-300 pA amplitude and 150 ms duration in cells initially maintained at their resting membrane potential (*V_rest_*), minimising the bridge potential that was not corrected in this analysis. Figure 3A shows an example of a recording from a *Scn2a^+/+^* cell at P31 to illustrate the variables that were analysed. Since the series resistance of the patch pipette in our experiments was in the 5-10 MΩ range, the error caused by the bridge potential in the estimate of the AP threshold (*AP_thr_*) could be as large as 3 mV. Thus, we focussed the analysis on two variables that are not affected by the bridge potential, namely the AP amplitude (*AP_amp_*), defined as the difference between the AP peak (*AP_peak_*) and *AP_thr_*, and of the AP width (*AP_wid_*), defined as the interval at which V_m_ is above the value corresponding to half of *AP_amp_*. The scatter plots of analysed somatic potential variables against age are reported in Supplemental Figure 3. Cells were again divided in three groups by age (P21-40, P21-24 and P30-40). Averaged somatic APs for the three strains and the three age groups are shown in Figure 3B. In these experiments, the largest difference could be qualitatively observed at P21-24 where the AP in *Scn2a^+/-^* cells appears wider and the AP in *Scn2a^+/L1314P^* cells appears smaller. The statistics (mean ± SD) of the variables for the three strains and the three age groups are reported in Figure 3C and Supplemental Table 4.

**Figure 3.**
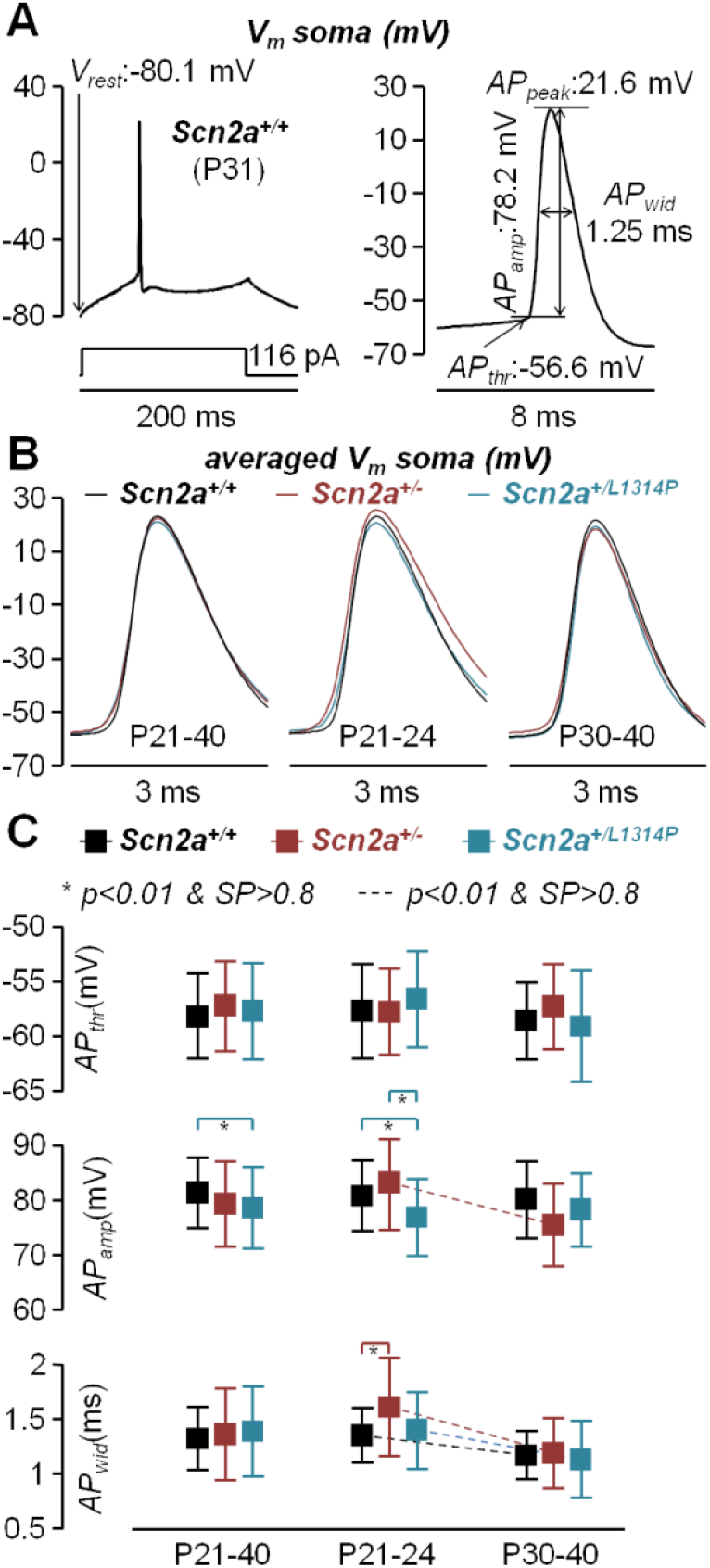
Somatic AP of *Scn2a^+/+^, Scn2a^+/-^* and *Scn2a^+/L1314P^* L5 pyramidal neurons. (A) On the left, somatic AP elicited by a 116 pA current pulse of 150 ms duration from a resting potential of -80.1 mV in a *Scn2a^+/+^*L5 pyramidal neuron at P31. On the right, illustration of the variables used for the analysis: threshold (*AP_thr_*, -56.6 mV), amplitude (AP_amp_, 78.2 mV) calculated as difference from *AP_thr_* and peak (*AP_peak_*, 21.6 mV), and width (*AP_wid_*, 1.25 ms) calculated as the interval at which V_m_ is above the value corresponding to half of *AP_amp_*. (B) Average of APs of all measured *Scn2a^+/+^* cells, *Scn2a^+/-^* cells, and *Scn2a^+/L1314P^* cells at P21-40, P21-24 and P30-40. Notice the greatest differences at P21-24. (C) Mean ± SD of *AP_thr_* (top), *AP_amp_* (middle) and *AP_wid_*(bottom) for *Scn2a^+/+^* cells, *Scn2a^+/-^* cells, and *Scn2a^+/L1314P^* cells at P21-40, P21-24 and P30-40. “*” indicates that the result of the two-sample ks-test between two groups of same age and different strain gave P<0.01 and SP>0.8. The dashed line indicates that the result of the two-sample ks-test between two age groups of the same strain gave P<0.01 and SP>0.8.

We assessed differences in the variables by applying the ks-test, using the P <0.01 / SP >0.8 criterion, and the results are reported in Supplemental Table 5. *AP_amp_* values in *Scn2a^+/L1314P^* cells were smaller than those in *Scn2a^+/+^* cells both at P21-40 (P=0.001093; SP=0.929) and in the youngest group (P=0.004413; SP=0.843). In younger *Scn2a^+/-^* cells, *AP_amp_* values were also larger than those in *Scn2a^+/L1314P^* cells (P=3.9·10^-5^; SP=0.989) and decreased in the *Scn2a^+/-^* oldest group (P=7.1·10^-5^; SP=0.996). *AP_wid_* values were larger in the young groups, with respect to the old groups, in *Scn2a^+/+^* cells (P=0.003498; SP=0.965), in *Scn2a^+/-^* cells (P=0.7·10^-5^; SP=0.999) and in *Scn2a^+/L1314P^* cells (P=8.0·10^-5^; SP=0.962). In the young group, *AP_wid_* values were larger in *Scn2a^+/-^* cells with respect to *Scn2a^+/+^* cells (P=0.000693; SP=0.947). These results cannot be explained in terms of VGNC changes only. A Na^+^ current increase leads to an increase of both *AP_amp_*and *AP_wid_*, but these two variables do not positively correlate. In contrast, a K^+^ current increase leads to a decrease of both *AP_amp_* and *AP_wid_*. Thus, the decrease with age of *AP_wid_* can be explained only by a K^+^ current increase. For the same reason, in young groups, the larger *AP_wid_* observed in *Scn2a^+/-^* cells and the smaller *AP_amp_* observed in *Scn2a^+/L1314P^* cells can be explained by concomitant smaller K^+^ current in the first case, and larger K^+^ current in the second case, with respect to *Scn2a^+/+^* cells. Somatic APs also contain information on AIS excitability. Thus we calculated the time derivative of the somatic V_m_ (dV_m_/dt) to extrapolate phase plots for the three strains, averaged over all ages (P21-40, Supplemental Figure 4A). Visually, these plots show that excitability is lower in both *Scn2a^+/-^*and *Scn2a^+/L1314P^* cells with respect to *Scn2a^+/+^* cells, with the lowest excitability observed in *Scn2a^+/L1314P^* cells. In all strains, dV_m_/dt peaks were between -30 mV and -20 mV and their values in individual cells are reported in the scatter plot of Supplemental Figure 4B. Notably, applying the ks-test on the dV_m_/dt peak over the ensemble of 142 *Scn2a^+/+^* cells and 136 *Scn2a^+/L1314P^* cells, gave P = 0.0033 and SP = 0.848. Similarly, in the analysis of the positive area of the phase plot (Supplemental Figure 4C), the ks-test gave P = 0.0014 and SP = 0.880, consistently with the hypothesis that *Scn2a^+/L1314P^* cells are less excitable because they express less functional Na_v_1.2 channels in the AIS. Different K^+^ currents are also expected to change the AP shape in the AIS, an aspect that we could directly investigate by V_m_ imaging.

### AIS APs in Scn2a^+/-^ and Scn2a^+/L1314P^ mice

Next, we investigated the kinetics of APs in the AIS. To this purpose, we loaded neurons with the voltage sensitive dye JPW1114 using a procedure described in several other reports (see for example [32,34]) and we recorded V_m_ signals associated with APs elicited by somatic current injection. Figure 4A shows schematically the three AIS positions (*d*, *m* and *p*) from where fluorescence was averaged and an example of recording from *Scn2a^+/+^* cell at P26. To analyse the kinetics of the AIS V_m_ waveforms, we applied to the ΔF/F_0_ traces a Savitzky-Golay (S-G) smoothing filter and defined the AIS *AP_wid_*(different from that defined in the soma) as the time interval during which V_m_ > 0.5 in the normalised AP (Figure 4B). As shown in this example, the AP becomes sharper as the distance from the soma increases, resulting in a smaller value of *AP_wid_* in the *d* position. The three examples of Figure 4C illustrate AIS V_m_ optical recordings associated with AP in a *Scn2a^+/+^* cell, in a *Scn2a^+/-^* cell and in a *Scn2a^+/L1314P^* cell at P23. In the *Scn2a^+/L1314P^* cell, the AP in *d* and *m* positions was visually sharper with respect to the other two cells. Altogether, we performed measurements 47 *Scn2a^+/+^* cells, 47 *Scn2a^+/-^* cells and 51 *Scn2a^+/L1314P^* cells, mostly in the two age groups P21-24 and P30-40. The histograms of *AP_wid_* in the three different positions for all groups are reported in Supplemental Figure 5. As for the other variables, we applied the ks-test with strong SP and high confidence level and we found that *AP_wid_*in *d* positions were shorter in *Scn2a^+/L1314P^* cells with respect to the other two strains at P21-24 and at P21-40 (Figure 4D). In addition, in both age groups, *AP_wid_* in *m* positions were shorter in *Scn2a^+/L1314P^* cells with respect to *Scn2a^+/-^* cells. The sharper distal APs in the AIS of *Scn2a^+/L1314P^* neurons match in the age groups with the smaller amplitudes of the somatic APs that we observed in the same strain (Figure 3C). Both results are consistent with the general hypothesis that larger K^+^ currents were activated in *Scn2a^+/L1314P^* cells.

**Figure 4.**
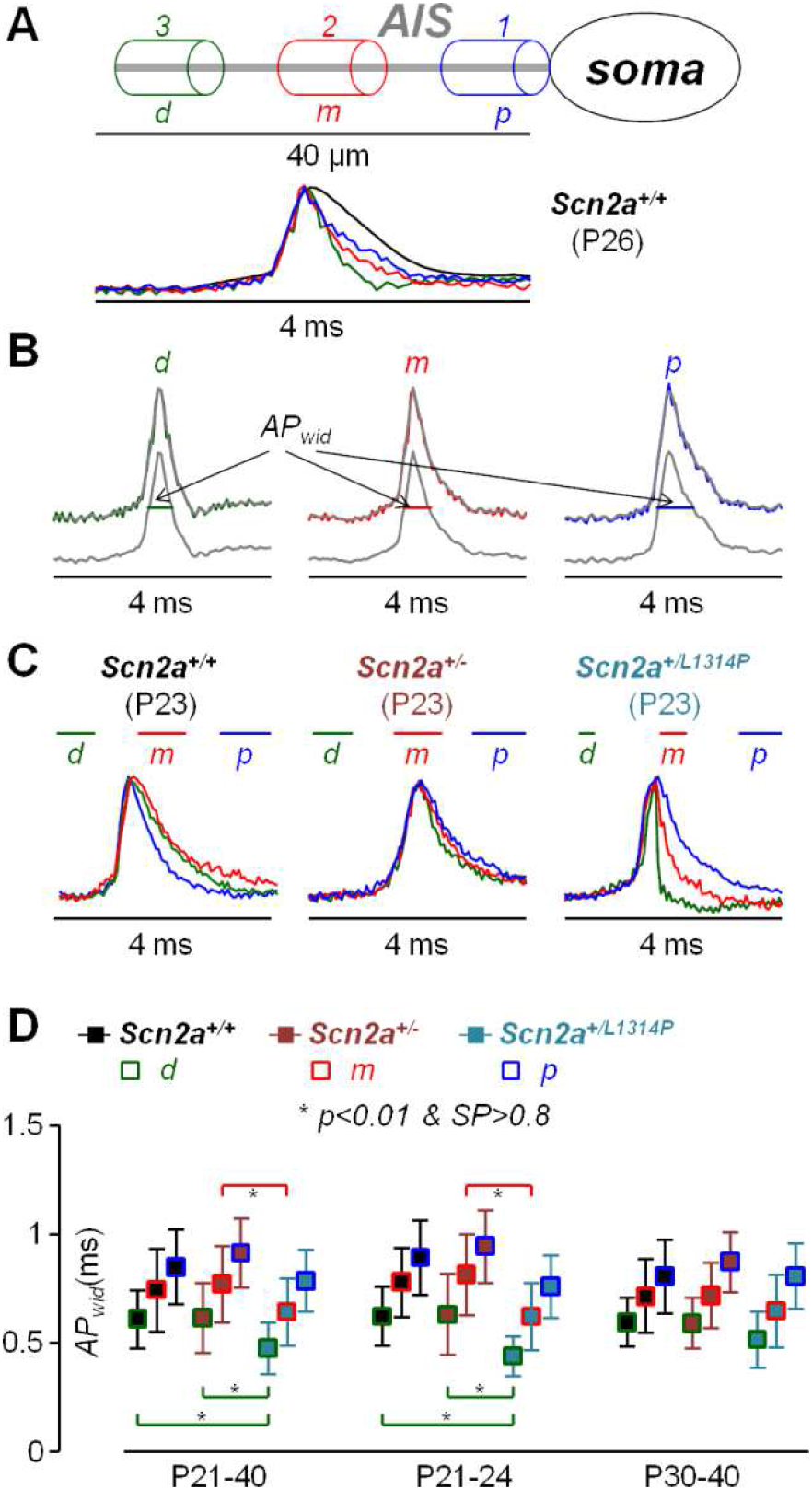
AIS APs in *Scn2a^+/+^, Scn2a^+/-^* and *Scn2a^+/L1314P^* L5 pyramidal neurons. (A) On the top, position of distal (*d*, green cylinder) medial (*m*, red cylinder) and proximal (*p*, blue cylinder) sites. On the bottom, from a *Scn2a^+/+^* L5 pyramidal neuron at P26, an AP in the soma (black trace) and in the three AIS sites (coloured traces) normalised and superimposed. (B) AIS traces in panel A smoothed with S-G filter and normalised to peak; *AP_wid_* is calculated as time for which V_m_ > 0.5. (C) AIS V_m_ optical recordings associated with AP in a *Scn2a^+/+^*cell, in a *Scn2a^+/-^* cell and in a *Scn2a^+/L1314P^* cell at P23. Coloured segments indicate *AP_wid_* in *d*, *m* and *p* positions. (D) Mean ± SD of *AP_wid_* in *d*, *m* and *p* positions for the three strains and the three age groups. Values used for these statistics are reported in Supplemental Figure 5. “*” indicates that the result of the two-sample ks-test between two groups gave P<0.01 and SP>0.8.

The AIS expresses several types of VGKCs and K_v_1 [35] and K_v_7 [36] are known to contribute to AP repolarisation. To explore the effect of inhibiting these two channels in the three strains, we delivered either 500 nM of the K_v_1 blocker dendrotoxin-K (dtx-K, [37]) or 10 µM of the K_v_7 blocker 10,10-*bis*(4-Pyridinylmethyl)-9(10*H*)-anthracenone dihydrochloride (xe991, [38]). Furthermore, we also tested the effect of delivery of 1 µM of the Ca^2+^-activated K^+^ channels (CAKCs) BK CAKCs inhibitor iberiotoxin (ibtx), since this channel contributes to AP kinetics in the AIS [28]. Figure 5A shows normalised APs in *d*, *m* and *p* positions from cells in the P21-24 group for each strain, before and after delivery of dtx-K. Blocking K_v_1 widened the AP at all AIS positions in every cell and the effect was larger where the AP was sharper at *d* positions. Figure 5B shows normalised APs in cells of the P21-24 group, in this case before and after delivery of xe991. Blocking K_v_7 also widened the AP at all AIS positions in every cell, but the effect did not appear larger in *d* positions. Finally, Figure 5C shows normalised APs before and after delivery of ibtx with widening caused by blocking BK channels similar to that caused by blocking K_v_7 channels. We tested the effect of delivery of each drug in 8-10 cells, belonging either to the P21-24 group or to the P30-40 group, in each of the three strains. The scatter plots of *AP_wid_* values at *d*, *m* and *p* positions before and after delivery of the drug, obtained in these cells, are reported in Supplemental Figure 6. Delivery of dtx-K, xe991 or ibtx either increased or did not change *AP_wid_* ratios. Figure 5D reports the mean ± SD of *AP_wid_* ratios in the three positions after and before delivery of each channel blocker (*AP_wid_^drug/control^*) in each strain. Application of paired t-test to *AP_wid_* values before and after drug delivery gave P values <0.01 in most positions of each strain and it was <0.05 otherwise, indicating that all channels (K_v_1, K_v_7 and BK) contribute to AP repolarisation.

**Figure 5.**
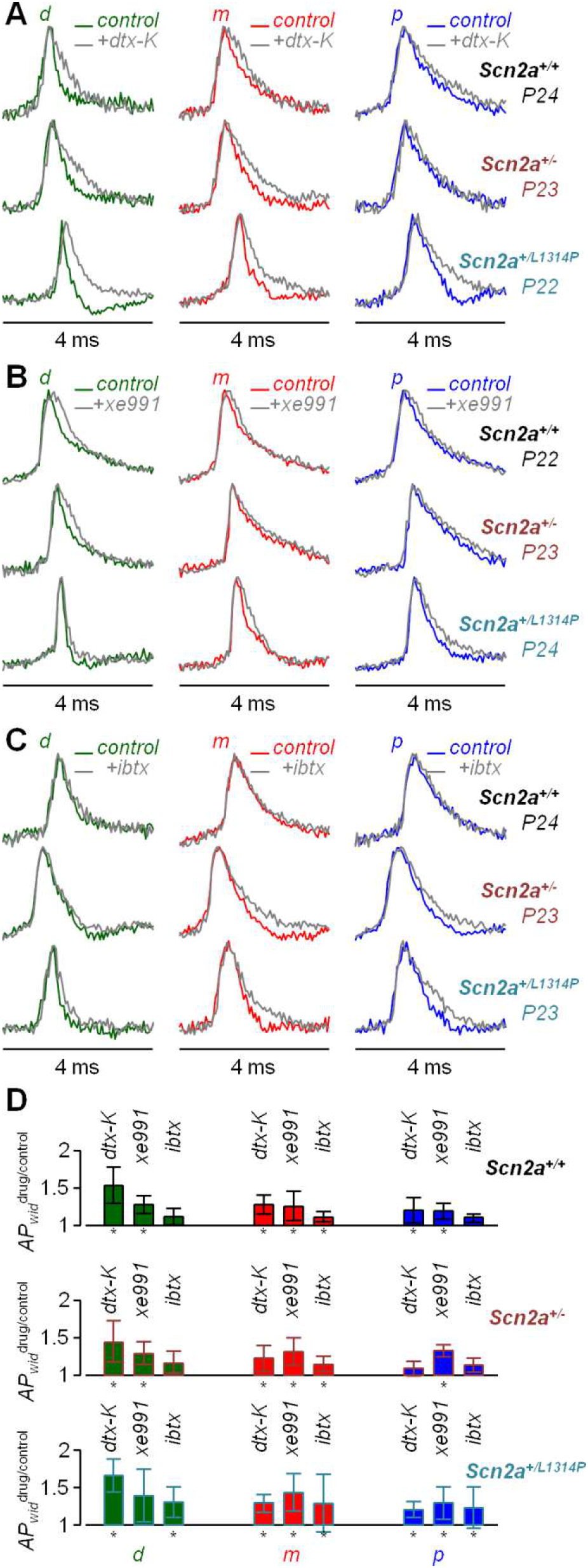
Effect of dtx-K, xe991 and ibtx on AIS APs in *Scn2a^+/+^, Scn2a^+/-^* and *Scn2a^+/L1314P^* L5 pyramidal neurons. (A) AIS V_m_ optical recordings associated with AP in a *Scn2a^+/+^*cell at P24, in a *Scn2a^+/-^* cell at P23 and in a *Scn2a^+/L1314P^* cell at P22 before (coloured traces) and after delivery of 500 nM dtx-K (K_v_1 inhibitor). (B) AIS V_m_ optical recordings associated with AP in a *Scn2a^+/+^*cell at P22, in a *Scn2a^+/-^* cell at P23 and in a *Scn2a^+/L1314P^* cell at P24 before (coloured traces) and after delivery of 10 µM xe991 (K_v_7 inhibitor). (C) AIS V_m_ optical recordings associated with AP in a *Scn2a^+/+^*cell at P24, in a *Scn2a^+/-^* cell at P23 and in a *Scn2a^+/L1314P^* cell at P23 before (coloured traces) and after delivery of 1 µM ibtx (BK inhibitor). (D) Mean ± SD of ratios *AP_wid_* after delivery of a drug and *AP_wid_*under control solution in *d*, *m* and *p* positions for the three strains. Values used for these statistics are reported in Supplemental Figure 6. “*” indicates that the result of paired t-test between two groups gave P<0.01.

Since each VGKC inhibition modulates V_m_ kinetics affecting all voltage-gated ion channels, these results cannot be directly correlated to channel distributions. The effect of dtx-K, however, was stronger at *d* positions in all three strains, indicating that K_v_1 plays a major role in the AIS AP sharpening. For this reason, although all channels may change in AIS remodelling, we restricted to K_v_1 changes in a minimal biophysical model that we built to recapitulate the experimental results obtained in this AIS analysis of the three strains.

#### Interpretation of experimental observations in terms of a minimal biophysical model

The complex set of multi-variable results can be generally explained by remodelling of the AIS during the critical juvenile period, which appears different in the three strains. This may implicate both AIS morphology and distributions of all ion channels and related protein. The AP generation can be biophysically modelled by simulating the kinetics of all ion channels in the AIS triggered by a depolarising pulse delivered in the soma, but the number of parameters involved in a realistic model would greatly exceed experimental information. In addition, the large cell-to-cell experimental variability should be mirrored by variability of model parameters, which can be matched by multiple parameter sets. Thus, instead of aiming at a realistic model, we searched for a minimal model where the AIS AP shape is essentially matched by computer simulations and where, consistently with the pharmacological analysis reported in Figure 5, the repolarising phase of the AIS AP is strongly regulated by K_v_1. Supplemental Figure 7A shows a NEURON-simulated AP at putative *d*, *m* and *p* sites of the AIS under “control” conditions and after removing 90% K_v_1, leading to AP widening that resembles the one obtained experimentally. The model, modified from a previously published one [28], contains several ion channels (including, among the others, Na_v_1.6, K_v_7 and BK). Although all distributions are expected to change in remodelling, we used a first approximation consisting of changing Na_v_1.2, K_v_1 and VGCCs (Ca_v_) only, in order to recapitulate results obtained in the three strains at the two juvenile stages. The distribution of these primary channels for the traces reported in Supplemental Figure 7A are illustrated by the straight black lines in Figure 6A. As shown in Supplemental Table 6, in this initial simulation, *AP_wid_* variables at the three AIS are consistent with mean experimental values in P21-24 *Scn2a^+/+^* mice. *R_p/d_*, calculated from the time integral of the absolute *I_Na_* (Figure 6B), was also very close to the experimental mean in P21-24 *Scn2a^+/+^* mice. Finally, to obtain *delay*, a 0.2 ms average filter was applied to the time integral of the absolute *I_Ca_*, the result was sampled at 10 kHz and fitted with a four-sigmoid function and the time interval calculated, in this way reproducing the analysis performed with experimental data (Figure 6C). Also *delay* in this computer simulation (200 µs) was close to the experimental mean in P21-24 *Scn2a^+/+^* mice (Supplemental Table 6). From this initial parameter set, we mimicked *Scn2a^+/+^* maturation from P21-24 to P30-40 by slightly increasing and shifting towards the soma Na_v_1.2, by increasing K_v_1 to match the AP shortening and by decreasing Ca_v_ to decrease the slow I_Ca_ component (Figure 6A and Supplemental Figure 7B). The NEURON-simulated AP traces with this parameter set are reported in Figure 6D and Supplemental Figure 7C. As shown in Supplemental Table 6, *R_p/d_*, *delay* and *AP_wid_*variables, extracted from this simulation, are consistent with mean experimental values in P30-40 *Scn2a^+/+^* mice, providing a basic maturation paradigm to analyse the other two strains. To mimic *Scn2a^+/-^* at P21-24, from the initial parameter set, we assumed a 20% LoF (fewer channels) and parameter changes in the opposite direction with respect to *Scn2a^+/+^* maturation: shift of Na_v_1.2 away from the soma, decrease of K_v_1 and slight increase of Ca_v_. A maturation paradigm from this early scenario (shift of Na_v_1.2 towards the soma, increase of K_v_1 and decrease of Ca_v_), but with the same Na_v_1.2 LoF, was used to mimic *Scn2a^+/-^* at P30-40. *R_p/d_*, *delay* and *AP_wid_* values, extracted from these two simulations, are consistent with mean experimental values in *Scn2a^+/-^* mice at both juvenile stages (Supplemental Table 6). Thus, it appears that experimental results in the *Scn2a^+/-^* mouse are consistent with a combination of Na_v_1.2 reduction and AIS developmental delay. To mimic *Scn2a^+/L1314P^* at P21-24, from the initial parameter set, we instead assumed a 50% Na_v_1.2 LoF and slight increase of Ca_v_, but in this case combined with a Na_v_1.2 shift towards the soma and a strong K_v_1 increase. In this strain, maturation was mimicked by partial recovery from disruption towards the parameter set of *Scn2a^+/+^* at P30-40: Na_v_1.2 LoF but with the same profile, recovered Ca_v_, but only partially recovered K_v_1. *R_p/d_*, *delay* and *AP_wid_* values from these simulations are consistent with mean experimental values in *Scn2a^+/L1314P^* mice at both juvenile stages. Thus, it appears that in addition to Na_v_1.2 LoF, the *Scn2a^+/L1314P^*mouse is characterised by AIS disruption, occurring at earlier developmental stages and producing an abnormal increase of VGKCs. This remodelling is likely the cause of AP waveform change experimentally observed in the *Scn2a^+/L1314P^* mouse model.

**Figure 6.**
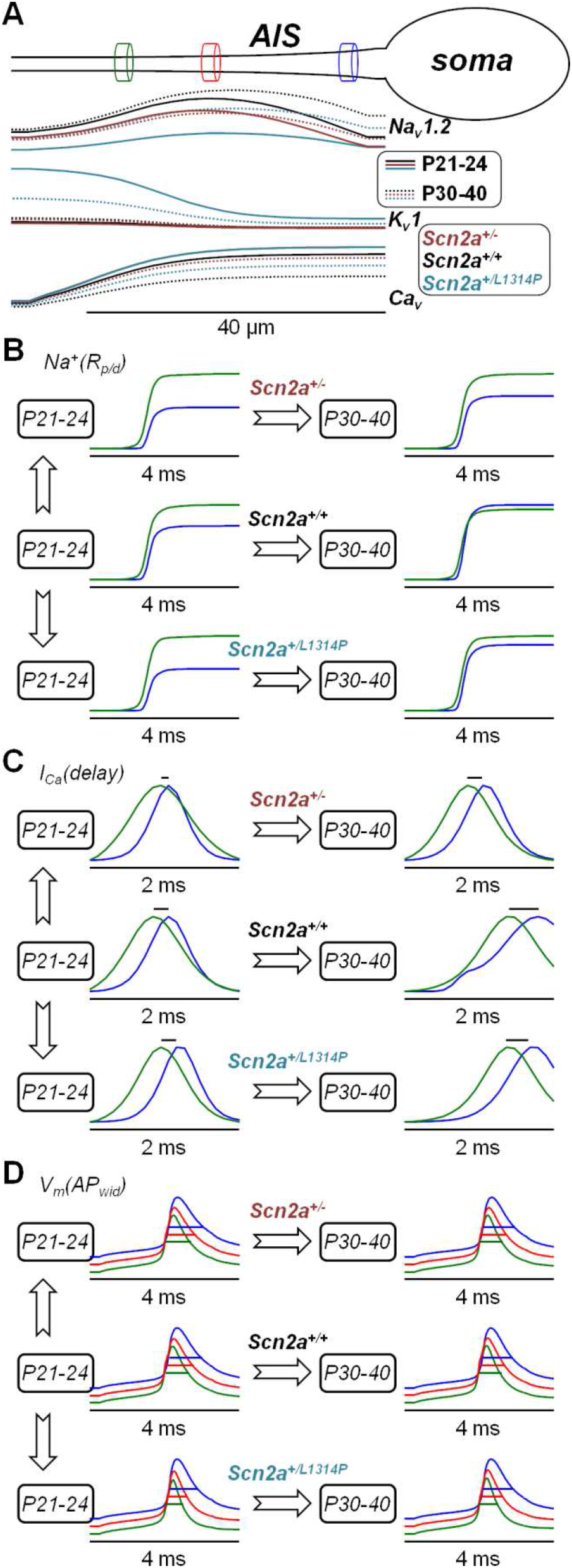
NEURON-simulations of “minimal” model mimicking juvenile AIS maturation in *Scn2a^+/+^, Scn2a^+/-^* and *Scn2a^+/L1314P^* L5 pyramidal neurons. (A) On the top, position of distal (*d*, green cylinder) medial (*m*, red cylinder) and proximal (*p*, blue cylinder) sites used to reproduce experimental results. On the bottom, profiles of Na_v_1.2, K_v_1 and Ca_v_ distributions in six simulations mimicking P21-24 (straight lines) to P30-40 maturation (dotted lined) in *Scn2a^+/+^, Scn2a^+/-^* and *Scn2a^+/L1314P^* L5 pyramidal neurons; percentage changes and shifts, with respect to the arrows, are reported in Supplemental Figure 7B. (B) Na^+^ influx, obtained from absolute *I_Na_* in the six simulations (reported in Supplemental Figure 7C) at *d* and *p* sites; values of R_p/d_ are reported in Supplemental Table 6. (C) From the six simulations, normalised *I_Ca_*, at *d* and *p* sites obtained after a procedure of filtering, integrating, re-sampling, fitting and calculating the time derivative, to reproduce the experimental measurement from the absolute *I_Ca_* traces reported in Supplemental Figure 7C; short black lines on the top of the traces show *delay* (values are reported in Supplemental Table 6). (D) From the six simulations, V_m_ at *d, m*, and *p* sites; short coloured lines in the middle of the traces show *AP_wid_* at each position (values are reported in Supplemental Table 6).

## Discussion

Haploinsufficiency of the *Scn2a* gene is considered a frequent cause of neurodevelopmental disorder characterized by ASD/ID. In haploinsufficient individuals, one *Scn2a* allele produces functional Na_v_1.2 channels, whereas the other is non-functional, leading to a reduction in overall Na_v_1.2 expression. Consequently, Na_v_1.2 LoF is generally considered the primary pathogenic mechanism underlying the associated phenotypes. Consistently, recent studies have shown that restoration of Na_v_1.2 expression through CRISPR-based transcriptional activation can ameliorate abnormalities in the *Scn2a*^+/-^ mouse model of *Scn2a* haploinsufficiency [39]. However, being implicated in brain development, Na_v_1.2 dysfunction is also expected to perturb structural and functional development in two possible concomitant ways: by plasticity mechanisms and/or by interaction with partner proteins. In the AIS, for instance, it was shown that the scaffold protein AnkG is regulated by Na_v_1.2 [40], affecting establishment and maintenance of the whole structure, including the distribution of other ion channels. This evidence raises the hypothesis that different LoF mutations of the *Scn2a* gene may not lead to the same developmental pathway. To our knowledge, the present study is the first one that addressed this hypothesis, by investigating AP generation in L5 pyramidal neurons in two distinct Na_v_1.2 LoF models. The two major findings of this study are: (1) that neurodevelopment of the AIS is different in *Scn2a^+/-^* and *Scn2a^+/L1314P^*mice; and (2) that functional and possibly structural secondary responses of the AIS, in particular involving K^+^ channels, are at least as important as Na_v_1.2 LoF.

### *Scn2a^+/-^* and *Scn2a^+/L1314P^* mouse models are different

The *Scn2a^+/-^* mouse has only one allele leading to Na_v_1.2 expression, whereas *Scn2a^+/L1314P^* DNA shares with the first model the same allele while the other one leads to expression of a Na_v_1.2 protein with a missense mutation. It was shown that the L1314P mutant *in vitro* is not targeted to the plasma membrane [30]. The amount of functional Na_v_1.2 transported to the AIS by AnkG and kinesin-1 [41] may not be linear with negative dominant effects induced by the L1314P mutant, but in both models a decrease of ion influx *via* Na_v_1.2 channels was expected at the beginning of this study. To address this question, we designed experiments on the basis of established evidence that Na_v_1.2 and Na_v_1.6 have different distribution in the AIS [12] and that Na_v_1.2 is permeable to Ca^2+^ [28,31]. Since Na_v_1.6 is mostly expressed in the distal part of the AIS, but Ca^2+^ influx *via* Na_v_1.2 is predominant at this site where APs are sharper, we expected a decrease in the variables *R_p/d_* and *delay* in both mouse models with respect to control mice. Instead, such a decrease was observed only in *Scn2a^+/-^*mice. A simple decrease of Na_v_1.2 in the AIS and the soma would translate into APs with smaller amplitude and, if reduced depolarisation decreases activation of K^+^ channels, AP widening. But results obtained from analysis of somatic APs were not consistent with this scenario. To address the problem, one must first consider that Na_v_1.2 alterations combine with developmental processes and that the time window of our analysis occurs in the juvenile phase of maturation. Classical electrophysiological studies performed in L5 pyramidal neurons of the visual [42] and prefrontal [43] cortex have shown that during the third postnatal week the somatic AP increases in amplitude and decreases in width, a phenomenon that can be attributed to the evolution of ion channel distribution in the AIS and in the soma. In the somato-sensory cortex, where the AIS length was measured using AnkG antibodies, it was found that the AIS elongates in the first two postnatal weeks and then it shortens in the following four weeks [44]. This evidence matches with our observation that the somatic *AP_wid_* decreases from the P21-24 group to the P30-40 group in all strains. Yet, with respect to *Scn2a^+/+^* cells, the somatic AP shape was found to be altered both in *Scn2a^+/-^* cells and in *Scn2a^+/L1314P^* cells, but in divergent manner. In the P21-24 group, somatic *AP_wid_*was larger in *Scn2a^+/-^* cells, whereas somatic *AP_amp_*was smaller in *Scn2a^+/L1314P^* cells when, concomitantly, the AP in the distal and medial part of the AIS was sharper. We infer that the two mouse models follow different developmental directions. In these paths, the impact of alteration of K^+^ distribution appears critical for neuronal excitability.

#### Impact of K^+^ channel function on AP generation in Na_v_1.2 LoF mouse models

In contrast to VGNCs and VGCCs, for which functional localisation can be explored by measuring Na^+^ and Ca^2+^ influx, investigation of K^+^ channels function relies on weaker information obtained from measurements of the AP waveform. Yet, the ability to image the AP along the AIS allowed basic exploration of some K^+^ channels types. Among those, K_v_1 is highly expressed in the AIS of L5 pyramidal neurons [45] and it plays a major role in AP repolarisation both in the AIS [35] and in axon collaterals [46]. K_v_7 regulates AP initiation and conduction [47] and knocking-out ankyrin-B, another important AIS scaffold protein, leads to down-regulation of K_v_7 [48]. Interestingly, in hippocampal pyramidal neurons, K_v_7 inhibition induces AIS plasticity [49]. Finally, CAKCs contribute to the regulation of AP waveforms [50], in particular during repolarisation, and BK channels are particularly efficiently activated by local Ca^2+^ influx within nanodomains [51]. Notably, in L5 pyramidal neurons dendrites they are selectively activated by N-type VGCCs Ca^2+^ influx [52], whereas in the AIS they are activated by Na_v_1.2 Ca^2+^ influx [28], making them an interesting candidate in regulation of K^+^ channels. Not surprisingly, selective block of each K^+^ channel type widened the AIS AP everywhere and in all strains, but the effect was more marked for K_v_1. This result guided the construction of a minimal biophysical model aimed at providing a computational framework to interpret the complex set of results of this study. This model suggests what can broadly happen in *Scn2a^+/-^* cells and in *Scn2a^+/L1314P^* cells. In *Scn2a^+/-^* cells, decrease in Na_v_1.2 expression appears combined with slower development. This combination leads to wider APs and reduction of *R_p/d_*and *delay* variables. In contrast, In *Scn2a^+/L1314P^* cells, enhanced clustering of K^+^ channels sharpens APs in the AIS, in combination with potentially larger decrease of Na_v_1.2 current because of negative dominant effects. The resulting AP sharpening limits Ca^2+^ entry via VGCCs and Na^+^ influx through Na_V_1.6 channels, and leading to values of *R_p/d_* and *delay* that are closer to wild-type values.

#### Significance of this study for future therapeutics

VGNCs, in particular Na_v_1.2 and Na_v_1.6, remain very promising therapeutic targets [53]. In particular, rescuing VGNC down-regulation in genetic disease [39] seems the most effective way for counteracting the underlying functional deficit. In the heterogeneity of ASD/ID [54], diagnosis occurs relatively late in development and the implementation of large-scale genetic screening before the onset of neurological symptoms [55] remains challenging in routine clinical practice. The present study highlights the importance of considering the temporal window of therapeutic intervention. Although two distinct *Scn2a* LoF mouse models display markedly different alterations in neuronal excitability and AIS maturation during the juvenile period, when pathological phenotypes are most evident, both exhibit partial spontaneous behavioural recovery in adulthood [14,29,56]. These findings suggest that different Na_v_1.2 LoF variants may drive distinct developmental trajectories in affected children. Thus, reversibility of ASD/ID pathophysiology as a function of disease stage and age at diagnosis should be a major focus of future therapeutic approaches.

## Methods

### Ethical approval and animal breeding

Experiments using animals were performed at the Laboratory of Interdisciplinary Physics in Grenoble in accordance with European Directives 2010/63/UE on laboratory animal use and care. Although below the minimal degree of severity, procedures were reviewed by the local ethics committee Cometh 12 affiliated to the animal facility of the University (# E3842110001). Mice were produced as follows: C57BL/6j females (*Scn2a^+/+^*), purchased from Charles River Laboratories (Saint-Germain-sur-l’Arbresle, France), were crossed with either *Scn2a^+/-^*males or *Scn2a^+/L1314P^* males (originally from IPMC, Valbonne-Sophia Antipolis, France). New-born mice were genotyped during the first week at the Plateforme de Génotypage, Neurocentre Magendie (Bordeaux, France), following guidelines from Jackson Laboratory. Females of all strains were used for experiments whereas *Scn2a^+/-^* males or *Scn2a^+/L1314P^*males were used for further breeding with new *Scn2a^+/+^* females. The age of crossing was 3-9 months and experiments in this report include animals from four generations. The choice of performing experiments on one gender (female) only was taken to restrict our analysis for a feasible number of animals in the project.

### Brain slice solution, preparation and maintenance

Female mice from all strains (21-40 postnatal days old) were anesthetised by isoflurane inhalation and the entire brain was removed after decapitation. The extracellular solution used for slice preparation maintenance and recordings contained (in mM): 125 NaCl, 26 NaHCO_3_, 1 MgSO_4_, 3 KCl, 1 NaH_2_PO_4_, 2 CaCl_2_ and 20 glucose, bubbled with 95% O_2_ and 5% CO_2_. Brain slices (350 µm thick) were prepared using the procedure described in earlier reports [27,28], using a Leica VT1200 vibratome (Wetzlar, Germany). Slices were incubated at 37°C for 45 minutes and maintained at room temperature before use.

### Electrophysiology

Experiments were performed using two electrophysiology setups: setup1: an upright Scientifica SliceScope microscope equipped with a motorised XY translation stage and PatchStar manipulators (Scientifica, Uckfield, UK), and a 60X Olympus water immersion objective (NA = 1); setup2 an Olympus BX51 mounted on a manual translational stage using a 60X NIKON water immersion objective (NA = 1). In both setups, patch-clamp recordings were made at the temperature of 32-34°C using Multiclamp 700A (Molecular Devices, Sunnyvale, CA) amplifiers. Slices with L5 pyramidal neurons in the somato-sensory cortex having an axon parallel to the slice surface were transferred to the recording chamber. The intracellular solution contained (in mM): 125 KMeSO_4_, 5 KCl, 8 MgSO_4_, 5 Na_2_-ATP, 0.3 Tris-GTP, 12 Tris-Phosphocreatine, 20 HEPES, adjusted to pH 7.35 with KOH. Electrical signals were acquired at 20 kHz and somatic APs were elicited in current clamp mode in two ways. For all imaging experiments, short (2-4 ms) current pulses of 1.5-3 nA were injected through the patch pipette. For recordings of APs only, current pulses of 100-300 pA were injected through the patch pipette.

### Imaging

Imaging methods were essentially the same utilised in a previous study [28]. All dyes were added to the intracellular solution and injected into cells. In Na^+^ imaging experiments, the Na^+^ indicator ING-2 (IonBiosciences, San Marcos, TX) was used at 0.5 mM and recordings were started 20 minutes after achieving the whole cell configuration. In Ca^2+^ imaging experiments, the Ca^2+^ indicator OG5N (Thermo Fisher Scientific) was used at 2 mM and recordings were started 20 minutes after achieving the whole cell configuration. In V_m_ imaging experiments, cell membranes were loaded with the voltage-sensitive dye D-2-ANEPEQ (JPW1114, 0.2 mg/mL, Thermo Fisher Scientific) for 30 minutes using a first patch clamp recording and then re-patched a second time with dye free solution. Na^+^ and Ca^2+^ imaging experiments and some V_m_ imaging experiments were performed in setup1, equipped with a LaserBank (Cairn Research, Faversham, UK). The laser beam was focussed onto the specimen using a custom-made telescope to obtain an illumination spot of ∼30 µm. Na^+^ and V_m_ indicators were excited using the 520 nm line band-pass filtered at 517 ± 10 nm and directed to the preparation using a 538 nm long-pass dichroic mirror. The Ca^2+^ indicator was excited using the 465 nm line band-pass filtered at 469 ± 17 nm and directed to the preparation using a 495 nm long-pass dichroic mirror. Na^+^ and V_m_ fluorescence emission signals were band-pass filtered at 559 ± 17 nm or long-pass filtered at >610 nm respectively. Ca^2+^ fluorescence emission was band-pass filtered at 525 ± 25 nm. Image sequences were demagnified by 0.5X and acquired with a DaVinci 2K CMOS camera (SciMeasure, Decatur, GA) with a pixel resolution of 500 nm, at 10 kHz in Na^+^ and Ca^2+^ imaging, or at 20 kHz in V_m_ imaging. The other V_m_ imaging experiments were performed in setup2 equipped with an LDI-7 laser (89 North, Williston, VT). Fluorescence was excited at 530 nm, band-pass filtered at 531 ± 40 nm and directed to the preparation using a 575 nm long-pass dichroic mirror. Fluorescence emission was long-pass filtered at >610 nm and acquired at 20 kHz either with a NeuroCCD camera (Redshirt Imaging, Decatur, GA) or with a Kinetix CMOS camera (Teledyne Photometrics, Tucson, AZ). In the first case, images were demagnified by 0.5 to obtain a pixel resolution of ∼2 µm, whereas in the second case, images were demagnified by 0.25 to obtain a pixel resolution of ∼433 nm.

#### Pharmacology

The three molecules used in this study were initially dissolved in water in stock solutions and small aliquots were kept at -20 °C before use. Dendrotoxin-K was purchased from Sigma-Aldrich, xe991 was purchased from Hello Bio (Dunshaughlin, Republic of Ireland) and iberiotoxin was purchased from Smartox Biotechnology (Saint Egrève, France). Molecules, diluted in extracellular solution at final concentrations indicated in the Results were delivered to AIS using a procedure described in a previous report [28].

#### Data analysis

Data analysis was performed on single trials in recordings of somatic AP only, and on averages of 4 trials with identical somatic response in imaging recordings. The analysis was performed using custom-written code in MATLAB. Somatic V_m_, measured with the patch electrode, was corrected for -11 mV junction potential. In shown somatic APs, elicited by 1.5-3 nA pulses (imaging experiments), the bridge was corrected offline by using the recorded injected current. These patch clamp recordings, where the bridge exceeded always 10 mV, were not quantitatively analysed. The somatic AP was analysed when elicited by longer (150 ms) current pulses of 100-300 pA and, in this case, the bridge was not corrected. This bridge, although smaller, biased the estimate of the real V_m_. We therefore estimated *AP_thr_* using the classical definition as the point at which the rate of rise of potential exceeds 50 V/sec [57], but we based our analysis on the bridge-independent variable *AP_AMP_* which is the difference between the AP peak and *AP_thr_*. The other variable extracted in analysis of somatic recordings was the AP width (*AP_wid_*), defined as the interval at which V_m_ is above the value corresponding to half of *AP_amp_*. In camera recordings, mean fluorescence from *d*, *m* or *p* sites (5 µm regions) was initially corrected for photo-bleaching using multi-exponential fits of fluorescence transients in single trials without evoked APs. Then, ΔF/F_0_ was calculated. For Na^+^ imaging experiments, ΔF/F_0_ signals were fitted with a model function described in an earlier report [27]. Since an absolute calibration of ING-2 is possible, ΔF/F_0_ was converted into Δ[Na^+^] and then into a positive charge superficial density using a general model of axon diameter. From this estimate, the Na^+^ current was extracted by calculating the time derivative (see [27]). Because this quantitative approach contains a potential margin of error and bias due to AIS morphological maturation, the comparative analysis was restricted to the ΔF/F_0_ only, where distances from the soma were the same regardless to AIS variability. For Ca^2+^ imaging experiments, the Ca^2+^ current was calculated as described in several previous reports [32,33] and the analysis focussed on delays between current peaks at different sites. Finally, for V_m_ imaging experiments, an absolute calibration of the AP size in the AIS was not available as in dendrites [34], and the noise fluctuations realistically corresponding to several mV prevented the analysis of amplitudes as in the soma. A S-G filter was applied to ΔF/F_0_ traces to smooth V_m_ signals and APs at each site were normalised to the peak. In contrast to somatic APs, since a measurement of the threshold cannot be achieved, the AIS *AP_wid_*at each site was defined as the time interval during which V_m_ > 0.5 in the normalised AP.

#### Statistical analysis

Statistical analysis performed to test a certain hypothesis was done in two distinct situations: 1) to assess whether a variable extracted from data analysis was the same in the three strains; 2) to assess whether delivery of a K^+^ channel inhibitor did not have any effect on a variable.

1. We adopted very stringent criteria on nonparametric tests to establish differences among the three strains. As a premise, in *Scn2a^+/+^* mice considered as control, variables changed from the P21-24 group to the P30-40 group. For this reason, statistical analysis comparing different strains was systematically conducted first within all analysed cells (P21-40) and then within cells belonging to the P21-24 and P30-40 groups. In Ca^2+^ imaging analysis, *delay* was essentially an integer multiplied by 100 µs. For this specific discrete variable, we used the two-sample Wilcoxon-Mann-Whitney (WMW) test. As this test does not directly offer an indication about effect magnitude, we complemented it with a standardized effect size (SES) that estimates of the strength of the observed effects and that was calculated as SES=Z/√N, where Z is the score of the test and N is the total number of observations [58]. Significant comparisons were those that met both P ˂ 0.01 and SES ˃ 0.5, where SES ˃ 0.5 represents a large effect size, so that reported findings reflect robust differences rather than trivial deviations. For all other variables that were not considered discrete, the ks-test was applied. This nonparametric test has the property to detect any distributional difference, including shifts in central tendency, variance, skewness, or the presence of subpopulations and it is therefore appropriate for this type of analysis. A caveat of this test, however, is its higher sensitivity near the centre of the distribution with respect to the tails, and its reduced statistical power for detecting small effect sizes. Therefore, to strengthen reliability, we implemented a dual criterion for statistical significance, that addresses both “false positive” and “false negative” errors. Results were considered significant only if P < 0.01, minimising false positives and ensuring that reported effects represent robust differences. But significance also required a statistical power higher than 0.8, where this threshold is generally accepted as appropriate cutoff for balancing between sensitivity and specificity [59]. SP for two samples was calculated using the standard Cohen’s formula [60]. No significance was reported when SP<0.8, even if the p-values were below 0.01.

2. In contrast to the first situation, analysis of the effect of K^+^ channel inhibitors was done in the same cell (APs before and after molecule delivery), allowing paired statistical testing. For this dataset, we performed a paired t-test comparing *AP_wid_* in control conditions and after local delivery of the K^+^ channel inhibitor. We considered P<0.01 as criterion to describe an effect as significant.

### Biophysical model and NEURON computation

All simulations were performed using the NEURON simulation environment (v.8.2.0). The model was based on a previously published biophysical model of mouse layer-5 pyramidal neurons [28,52], including neuronal morphology, passive membrane properties, temperature settings, ion channel kinetics and dendritic conductance distributions, which were left unchanged in the present study. Model and simulation files will be made available in the ModelDB section of the Senselab database (https://modeldb.science/2041700). The original model consists of a multicompartmental NEURON implementation of a reconstructed L5 pyramidal neuron, including an explicit axon initial segment (AIS) with a length of L=40 µm, and incorporates a full complement of voltage-gated Na*, K* and Ca²* conductances distributed according to experimental and theoretical constraints described in the original studies. Briefly, active properties include Na_v_1.2 and Na_v_1.6 sodium channels; multiple potassium conductance (delayed rectifier, K_v_1, K_v_7, SK and BK Ca²*-dependent channels); a non-specific I_h_ current; and several types of voltage-gated Ca²* channels (T-, L-, R-, N-, P/Q- type, low and high threshold), together with a simple Ca²* extrusion mechanism. In the present work, we focused specifically on the AIS and used the same ion-channel mechanisms introduced in a previous study [28] for this compartment, including Na_v_1.2, Na_v_1.6, K_v_1, K_v_7, and low- and high-threshold Ca_v_ currents, together with a new parameterized description of their spatial distributions, in accordance with experimental evidence [12,35]. BK Ca²*-dependent K* channels were also present in the AIS, with a linearly increasing conductance density from the soma toward the distal AIS [28], and were not modified in the present study. In addition, SK and delayed rectifier potassium channels were included in the AIS with a uniform conductance density and were likewise left unchanged. Rather than modifying channel kinetics, we controlled AIS excitability by reshaping the spatial profiles and total densities of selected ion channels along the AIS. To control the spatial profile of ion channel densities, we implemented a parameterized distribution function defined as the product of two sigmoidal functions. For each AIS mechanism, the local maximal conductance at a normalized position x∈[0,1] along the AIS was computed as

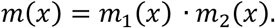

where the two sigmoidal components define the proximal and distal transitions:

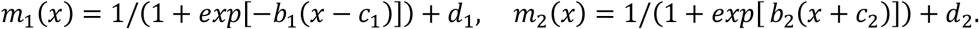

Here, the parameters (*b*_1_, *c*_1_, *d*_1_) control the slope, centre and offset of the proximal sigmoid, whereas (*b*_2_, *c*_2_, *d*_2_) define the corresponding properties of the distal sigmoid. Their product generates smooth and asymmetric profiles capable of producing monotonic, peaked or plateau-like distributions depending on parameter choice. To ensure that changes in the shape of the distribution did not alter the total conductance available along the AIS, each profile was normalised by its integral.

## Data and code availability

The complete dataset of imaging/electrophysiological experiments in brain slices, used in this study, is available in the public repository Zenodo (doi: 10.5281/zenodo.23809825).

## Acknowledgements

This work was funded by the *Agence Nationale de la Recherche* through two grants (ANR-21-CE18-0042 – Nav12RESCUE and Labex *Ion Channels Science and Therapeutics:* program number ANR-11-LABX-0015). We acknowledge the support of the Université Côte d’Azur’s Initiative of Excellence (IdEx) Jedi (ANR-15-IDEX-01). E.G. was funded by the Regione Sicilia (CUP G79J21012770001). We are indebted to the *Fédération pour la Recherche sur le Cerveau* (FRC – Grant *Espoir en tête*, Rotary France) for financing part of the experimental equipment. We acknowledge a contribution from the Italian National Recovery and Resilience Plan (NRRP), M4C2, funded by the European Union – NextGenerationEU (Project IR0000011, CUP B51E22000150006, “EBRAINS-Italy”). This project has also received funding from the EU’s Horizon Europe research and innovation program under grant agreement 101147319.

## Supplemental Tables

**Supplemental Table 1.** P and SP values from two-sample ks-tests performed to the Na^+^ influx parameter *R_p/d_* between various age groups of *Scn2a^+/+^*, *Scn2a^+/-^* and *Scn2a^+/L1314P^* cells. In bold characters the tests giving SP > 0.8 and P < 0.01.

| *GROUPS* | *P* | *Statistical Power (SP)* |
|---|---|---|
| *Scn2a*<sup>+/+</sup> P21-24 vs *Scn2a*<sup>+/+</sup> P30-40 | 0.007596 | 0.757 |
| *Scn2a*<sup>+/-</sup> P21-24 vs *Scn2a*<sup>+/-</sup> P30-40 | 0.014463 | 0.509823 |
| ***Scn2a*<sup>+/L1314P</sup> P21-24 vs *Scn2a*<sup>+/L1314P</sup> P30-40** | **0.003862** | **0.970** |
| ***Scn2a*<sup>+/+</sup> P21-40 vs *Scn2a*<sup>+/-</sup> P21-40** | **0.002335** | **0.952** |
| *Scn2a*<sup>+/+</sup> P21-24 vs *Scn2a*<sup>+/-</sup> P21-24 | 0.048117 | 0.757 |
| ***Scn2a*<sup>+/+</sup> P30-40 vs *Scn2a*<sup>+/-</sup> P30-40** | **0.007596** | **0.821** |
| *Scn2a*<sup>+/+</sup> P21-40 vs *Scn2a*<sup>+/L1314P</sup> P21-40 | 0.458 | 0.407124 |
| *Scn2a*<sup>+/+</sup> P21-24 vs *Scn2a*<sup>+/L1314P</sup> P21-24 | 0.155576 | 0.590 |
| *Scn2a*<sup>+/+</sup> P30-40 vs *Scn2a*<sup>+/L1314P</sup> P30-40 | 0.488908 | 0.253 |
| *Scn2a*<sup>+/-</sup> P21-40 vs *Scn2a*<sup>+/L1314P</sup> P21-40 | 0.680 | 0.120028 |
| *Scn2a*<sup>+/-</sup> P21-24 vs *Scn2a*<sup>+/L1314P</sup> P21-24 | 0.929326 | 0.074 |
| *Scn2a*<sup>+/-</sup> P30-40 vs *Scn2a*<sup>+/L1314P</sup> P30-40 | 0.028046 | 0.814 |

**Supplemental Table 2.** P and SES values from two-sample wmw-tests applied to “*delay*” in various age groups of *Scn2a^+/+^*, *Scn2a^+/-^* and *Scn2a^+/L1314P^* cells. In bold characters the tests giving SES > 0.5 and P < 0.01.

| *GROUPS* | *P* | ***Standardised Effect Size (SES)*** |
|---|---|---|
| ***Scn2a*<sup>+/+</sup> P21-24 vs *Scn2a*<sup>+/+</sup> P30-40** | **0.001164** | **0.66** |
| *Scn2a*<sup>+/-</sup> P21-24 vs *Scn2a*<sup>+/-</sup> P30-40 | 0.1709 | 0.23 |
| *Scn2a*<sup>+/L1314P</sup> P21-24 vs *Scn2a*<sup>+/L1314P</sup> P30-40 | 0.1268 | 0.29 |
| ***Scn2a*<sup>+/+</sup> P21-40 vs *Scn2a*<sup>+/-</sup> P21-40** | **0.000001253** | **0.55** |
| *Scn2a*<sup>+/+</sup> P21-24 vs *Scn2a*<sup>+/-</sup> P21-24 | 0.5338 | 0.12 |
| ***Scn2a*<sup>+/+</sup> P30-40 vs *Scn2a*<sup>+/-</sup> P30-40** | **0.00001094** | **0.75** |
| *Scn2a*<sup>+/+</sup> P21-40 vs *Scn2a*<sup>+/L1314P</sup> P21-40 | 0.3371 | 0.11 |
| *Scn2a*<sup>+/+</sup> P21-24 vs *Scn2a*<sup>+/L1314P</sup> P21-24 | 0.9100 | 0.23 |
| *Scn2a*<sup>+/+</sup> P30-40 vs *Scn2a*<sup>+/L1314P</sup> P30-40 | 0.08518 | 0.33 |
| *Scn2a*<sup>+/-</sup> P21-40 vs *Scn2a*<sup>+/L1314P</sup> P21-40 | 0.0002226 | 0.42 |
| *Scn2a*<sup>+/-</sup> P21-24 vs *Scn2a*<sup>+/L1314P</sup> P21-24 | 0.9902 | 0.19 |
| ***Scn2a*<sup>+/-</sup> P30-40 vs *Scn2a*<sup>+/L1314P</sup> P30-40** | **0.001769** | **0.51** |

**Supplemental Table 3.**
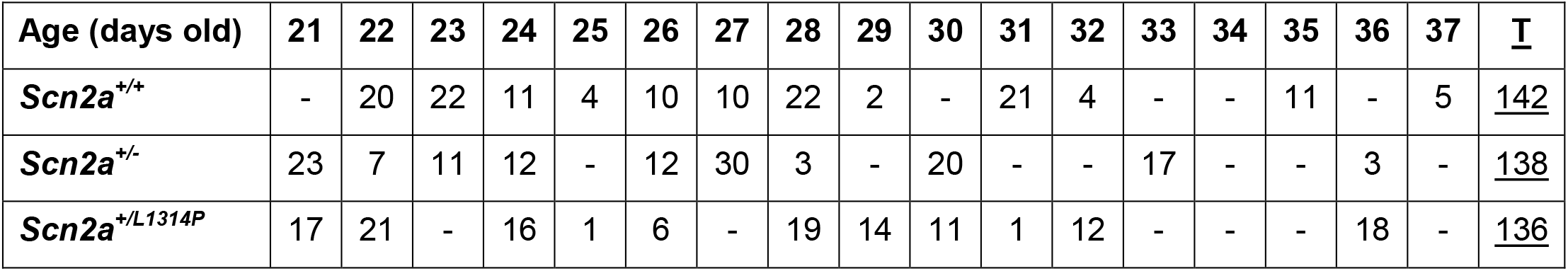
Summary of number of cells at different postnatal days analysed for somatic AP in 15 *Scn2a^+/+^* mice, 12 *Scn2a^+/-^* mice and 14 *Scn2a^+/L1314P^* mice.

| **Age (days old)** | **21** | **22** | **23** | **24** | **25** | **26** | **27** | **28** | **29** | **30** | **31** | **32** | **33** | **34** | **35** | **36** | **37** | **T** |
|---|---|---|---|---|---|---|---|---|---|---|---|---|---|---|---|---|---|
| ***Scn2a<sup>+/+</sup>*** | - | 20 | 22 | 11 | 4 | 10 | 10 | 22 | 2 | - | 21 | 4 | - | - | 11 | - | 5 | 142 |
| ***Scn2a<sup>+/-</sup>*** | 23 | 7 | 11 | 12 | - | 12 | 30 | 3 | - | 20 | - | - | 17 | - | - | 3 | - | 138 |
| ***Scn2a<sup>+/L1314P</sup>*** | 17 | 21 | - | 16 | 1 | 6 | - | 19 | 14 | 11 | 1 | 12 | - | - | - | 18 | - | 136 |

**Supplemental Table 4.** Mean ± SD of AP_thr_, AP_amp_ and AP_wid_ in cells from 15 *Scn2a^+/+^* female mice, 12 *Scn2a^+/-^* female mice and 14 *Scn2a^+/L1314P^*female mice at three time windows (P21-40, P21-24 and P30-40). The number of cells (N) for each population is indicated.

| *AP<sub>thr</sub>* (mV) | **P21-40** | **P21-24** | **P30-40** |
|---|---|---|---|
| ***Scn2a<sup>+/+</sup>*** | -58.2 ± 3.9 (N=142) | -57.7 ± 4.3 (N=53) | -58.6 ± 3.5 (N=41) |
| ***Scn2a<sup>+/-</sup>*** | -57.2 ± 4.1 (N=138) | -57.8 ± 3.9 (N=53) | -57.3 ± 3.9 (N=40) |
| ***Scn2a<sup>+/L1314P</sup>*** | -57.7 ± 4.4 (N=136) | -56.6 ± 4.4 (N=54) | --59.1 ± 5.1 (N=42) |
| *AP<sub>amp</sub>* (mV) | **P21-40** | **P21-24** | **P30-40** |
| ***Scn2a<sup>+/+</sup>*** | 81.4 ± 6.4 (N=142) | 80.8 ± 6.4 (N=53) | 80.2 ± 7.0 (N=41) |
| ***Scn2a<sup>+/-</sup>*** | 79.4 ± 7.9 (N=138) | 83.2 ± 8.3 (N=53) | 75.5 ± 7.5 (N=40) |
| ***Scn2a<sup>+/L1314P</sup>*** | 78.6 ± 7.4* (N=136) | 76.9 ± 7.0* (N=54) | 78.3 ± 6.7 (N=42) |
| *AP<sub>wid</sub>* (ms) | **P21-40** | **P21-24** | **P30-40** |
| ***Scn2a<sup>+/+</sup>*** | 1.32 ± 0.29 (N=142) | 1.35 ± 0.25 (N=53) | 1.17 ± 0.22 (N=41) |
| ***Scn2a<sup>+/-</sup>*** | 1.36 ± 0.42 (N=138) | 1.61 ± 0.45* (N=53) | 1.19 ± 0.32 (N=40) |
| ***Scn2a<sup>+/L1314P</sup>*** | 1.39 ± 0.41 (N=136) | 1.40 ± 0.34 (N=54) | 1.13 ± 0.35 (N=42) |

**Supplemental Table 5.**
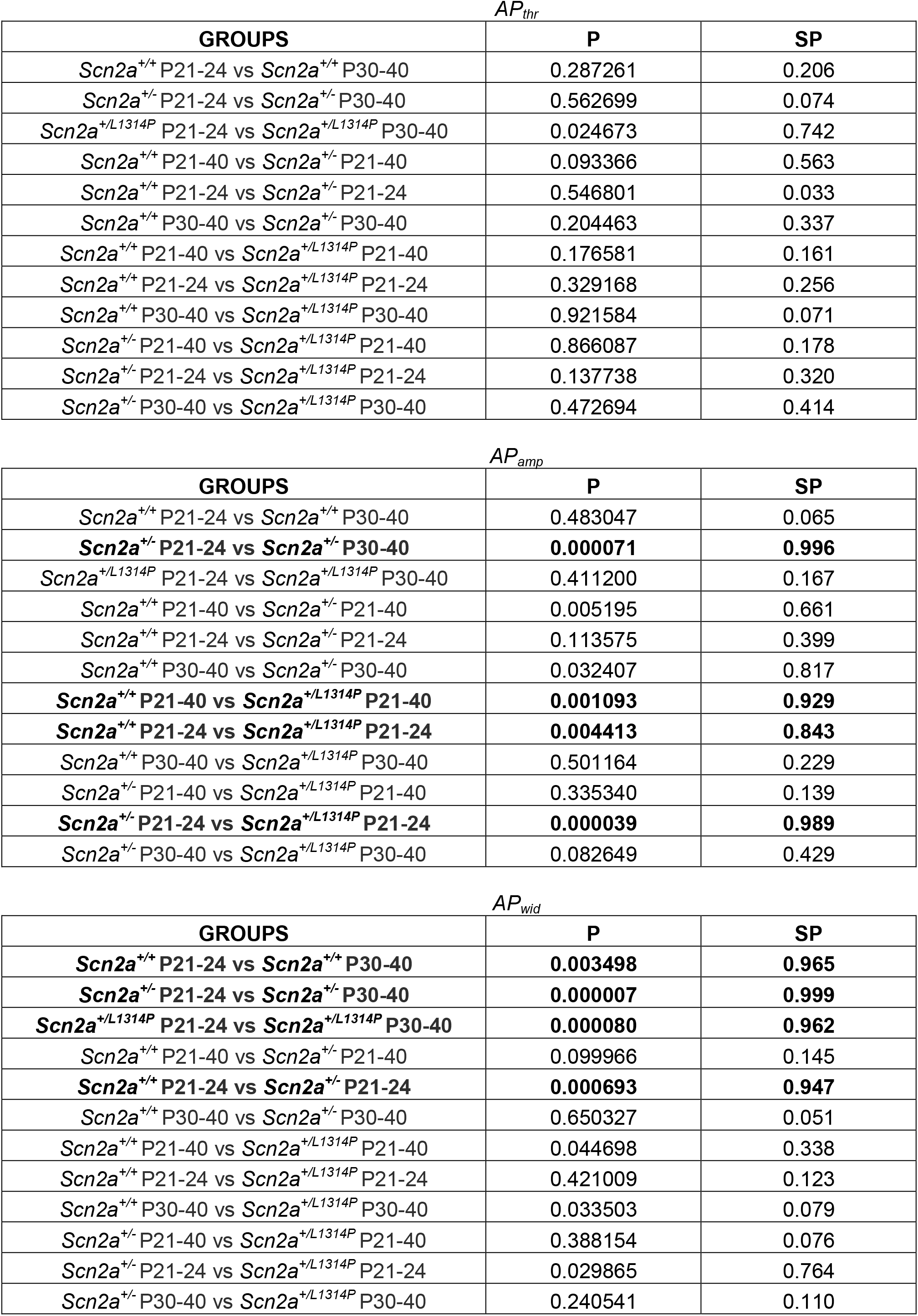
P and SP values from two-sample ks-tests performed to the somatic AP potential parameters *AP_thr_*, *AP_amp_* and *AP_wid_* between various age groups of *Scn2a^+/+^*, *Scn2a^+/-^* and *Scn2a^+/L1314P^* cells. In bold characters the tests giving SP > 0.8 and P < 0.01.

| *AP<sub>thr</sub>* |  |  |
|---|---|---|
| **GROUPS** | **P** | **SP** |
| *Scn2a<sup>+/+</sup>* P21-24 vs *Scn2a<sup>+/+</sup>* P30-40 | 0.287261 | 0.206 |
| *Scn2a<sup>+/-</sup>* P21-24 vs *Scn2a<sup>+/-</sup>* P30-40 | 0.562699 | 0.074 |
| *Scn2a<sup>+/L1314P</sup>* P21-24 vs *Scn2a<sup>+/L1314P</sup>* P30-40 | 0.024673 | 0.742 |
| *Scn2a<sup>+/+</sup>* P21-40 vs *Scn2a<sup>+/-</sup>* P21-40 | 0.093366 | 0.563 |
| *Scn2a<sup>+/+</sup>* P21-24 vs *Scn2a<sup>+/-</sup>* P21-24 | 0.546801 | 0.033 |
| *Scn2a<sup>+/+</sup>* P30-40 vs *Scn2a<sup>+/-</sup>* P30-40 | 0.204463 | 0.337 |
| *Scn2a<sup>+/+</sup>* P21-40 vs *Scn2a<sup>+/L1314P</sup>* P21-40 | 0.176581 | 0.161 |
| *Scn2a<sup>+/+</sup>* P21-24 vs *Scn2a<sup>+/L1314P</sup>* P21-24 | 0.329168 | 0.256 |
| *Scn2a<sup>+/+</sup>* P30-40 vs *Scn2a<sup>+/L1314P</sup>* P30-40 | 0.921584 | 0.071 |
| *Scn2a<sup>+/-</sup>* P21-40 vs *Scn2a<sup>+/L1314P</sup>* P21-40 | 0.866087 | 0.178 |
| *Scn2a<sup>+/-</sup>* P21-24 vs *Scn2a<sup>+/L1314P</sup>* P21-24 | 0.137738 | 0.320 |
| *Scn2a<sup>+/-</sup>* P30-40 vs *Scn2a<sup>+/L1314P</sup>* P30-40 | 0.472694 | 0.414 |
| *AP<sub>amp</sub>* |  |  |
| **GROUPS** | **P** | **SP** |
| *Scn2a<sup>+/+</sup>* P21-24 vs *Scn2a<sup>+/+</sup>* P30-40 | 0.483047 | 0.065 |
| ***Scn2a<sup>+/-</sup>*** **P21-24 vs** ***Scn2a<sup>+/-</sup>*** **P30-40** | **0.000071** | **0.996** |
| *Scn2a<sup>+/L1314P</sup>* P21-24 vs *Scn2a<sup>+/L1314P</sup>* P30-40 | 0.411200 | 0.167 |
| *Scn2a<sup>+/+</sup>* P21-40 vs *Scn2a<sup>+/-</sup>* P21-40 | 0.005195 | 0.661 |
| *Scn2a<sup>+/+</sup>* P21-24 vs *Scn2a<sup>+/-</sup>* P21-24 | 0.113575 | 0.399 |
| *Scn2a<sup>+/+</sup>* P30-40 vs *Scn2a<sup>+/-</sup>* P30-40 | 0.032407 | 0.817 |
| ***Scn2a<sup>+/+</sup>*** **P21-40 vs** ***Scn2a<sup>+/L1314P</sup>*** **P21-40** | **0.001093** | **0.929** |
| ***Scn2a<sup>+/+</sup>*** **P21-24 vs** ***Scn2a<sup>+/L1314P</sup>*** **P21-24** | **0.004413** | **0.843** |
| *Scn2a<sup>+/+</sup>* P30-40 vs *Scn2a<sup>+/L1314P</sup>* P30-40 | 0.501164 | 0.229 |
| *Scn2a<sup>+/-</sup>* P21-40 vs *Scn2a<sup>+/L1314P</sup>* P21-40 | 0.335340 | 0.139 |
| ***Scn2a<sup>+/-</sup>*** **P21-24 vs** ***Scn2a<sup>+/L1314P</sup>*** **P21-24** | **0.000039** | **0.989** |
| *Scn2a<sup>+/-</sup>* P30-40 vs *Scn2a<sup>+/L1314P</sup>* P30-40 | 0.082649 | 0.429 |
| *AP<sub>wid</sub>* |  |  |
| **GROUPS** | **P** | **SP** |
| ***Scn2a<sup>+/+</sup>*** **P21-24 vs** ***Scn2a<sup>+/+</sup>*** **P30-40** | **0.003498** | **0.965** |
| ***Scn2a<sup>+/-</sup>*** **P21-24 vs** ***Scn2a<sup>+/-</sup>*** **P30-40** | **0.000007** | **0.999** |
| ***Scn2a<sup>+/L1314P</sup>*** **P21-24 vs** ***Scn2a<sup>+/L1314P</sup>*** **P30-40** | **0.000080** | **0.962** |
| *Scn2a<sup>+/+</sup>* P21-40 vs *Scn2a<sup>+/-</sup>* P21-40 | 0.099966 | 0.145 |
| ***Scn2a<sup>+/+</sup>*** **P21-24 vs** ***Scn2a<sup>+/-</sup>*** **P21-24** | **0.000693** | **0.947** |
| *Scn2a<sup>+/+</sup>* P30-40 vs *Scn2a<sup>+/-</sup>* P30-40 | 0.650327 | 0.051 |
| *Scn2a<sup>+/+</sup>* P21-40 vs *Scn2a<sup>+/L1314P</sup>* P21-40 | 0.044698 | 0.338 |
| *Scn2a<sup>+/+</sup>* P21-24 vs *Scn2a<sup>+/L1314P</sup>* P21-24 | 0.421009 | 0.123 |
| *Scn2a<sup>+/+</sup>* P30-40 vs *Scn2a<sup>+/L1314P</sup>* P30-40 | 0.033503 | 0.079 |
| *Scn2a<sup>+/-</sup>* P21-40 vs *Scn2a<sup>+/L1314P</sup>* P21-40 | 0.388154 | 0.076 |
| *Scn2a<sup>+/-</sup>* P21-24 vs *Scn2a<sup>+/L1314P</sup>* P21-24 | 0.029865 | 0.764 |
| *Scn2a<sup>+/-</sup>* P30-40 vs *Scn2a<sup>+/L1314P</sup>* P30-40 | 0.240541 | 0.110 |

**Supplemental Table 6.** Comparison between *R_p/d_*, *delay* and *AP_wid_* values obtained experimentally (mean) in *Scn2a^+/+^*, *Scn2a^+/-^* and *Scn2a^+/L1314P^* mice and values extracted from NEURON-simulations reported in Figure 6B-D.

| ***R<sub>p/d</sub>*** |  | **P21-24** | **P30-40** |
|---|---|---|---|
| ***Scn2a<sup>+/+</sup>*** | Experiment (mean) | 0.73 | 1.12 |
|  | Simulation | 0.72 | 1.06 |
| ***Scn2a<sup>+/-</sup>*** | Experiment (mean) | 0.56 | 0.69 |
|  | Simulation | 0.54 | 0.70 |
| ***Scn2a<sup>+/L1314P</sup>*** | Experiment (mean) | 0.59 | 0.92 |
|  | Simulation | 0.56 | 0.88 |
| ***delay (µs)*** |  | **P21-24** | **P30-40** |
| ***Scn2a<sup>+/+</sup>*** | Experiment (mean) | 183 | 383 |
|  | Simulation | 200 | 400 |
| ***Scn2a<sup>+/-</sup>*** | Experiment (mean) | 105 | 146 |
|  | Simulation | 100 | 200 |
| ***Scn2a<sup>+/L1314P</sup>*** | Experiment (mean) | 185 | 273 |
|  | Simulation | 200 | 300 |
| ***proximal AP<sub>wid</sub>*** **(µs)** |  | **P21-24** | **P30-40** |
| ***Scn2a<sup>+/+</sup>*** | Experiment (mean) | 890 | 800 |
|  | Simulation | 875 | 800 |
| ***Scn2a<sup>+/-</sup>*** | Experiment (mean) | 940 | 870 |
|  | Simulation | 925 | 850 |
| ***Scn2a<sup>+/L1314P</sup>*** | Experiment (mean) | 810 | 760 |
|  | Simulation | 750 | 800 |
| ***medial AP<sub>wid</sub>*** **(µs)** |  | **P21-24** | **P30-40** |
| ***Scn2a<sup>+/+</sup>*** | Experiment (mean) | 780 | 710 |
|  | Simulation | 750 | 700 |
| ***Scn2a<sup>+/-</sup>*** | Experiment (mean) | 820 | 710 |
|  | Simulation | 825 | 700 |
| ***Scn2a<sup>+/L1314P</sup>*** | Experiment (mean) | 620 | 650 |
|  | Simulation | 550 | 625 |
| ***distal AP<sub>wid</sub>*** **(µs)** |  | **P21-24** | **P30-40** |
| ***Scn2a<sup>+/+</sup>*** | Experiment (mean) | 630 | 600 |
|  | Simulation | 675 | 600 |
| ***Scn2a<sup>+/-</sup>*** | Experiment (mean) | 630 | 590 |
|  | Simulation | 700 | 625 |
| ***Scn2a<sup>+/L1314P</sup>*** | Experiment (mean) | 430 | 520 |
|  | Simulation | 475 | 525 |

**Supplemental Figure 1.**
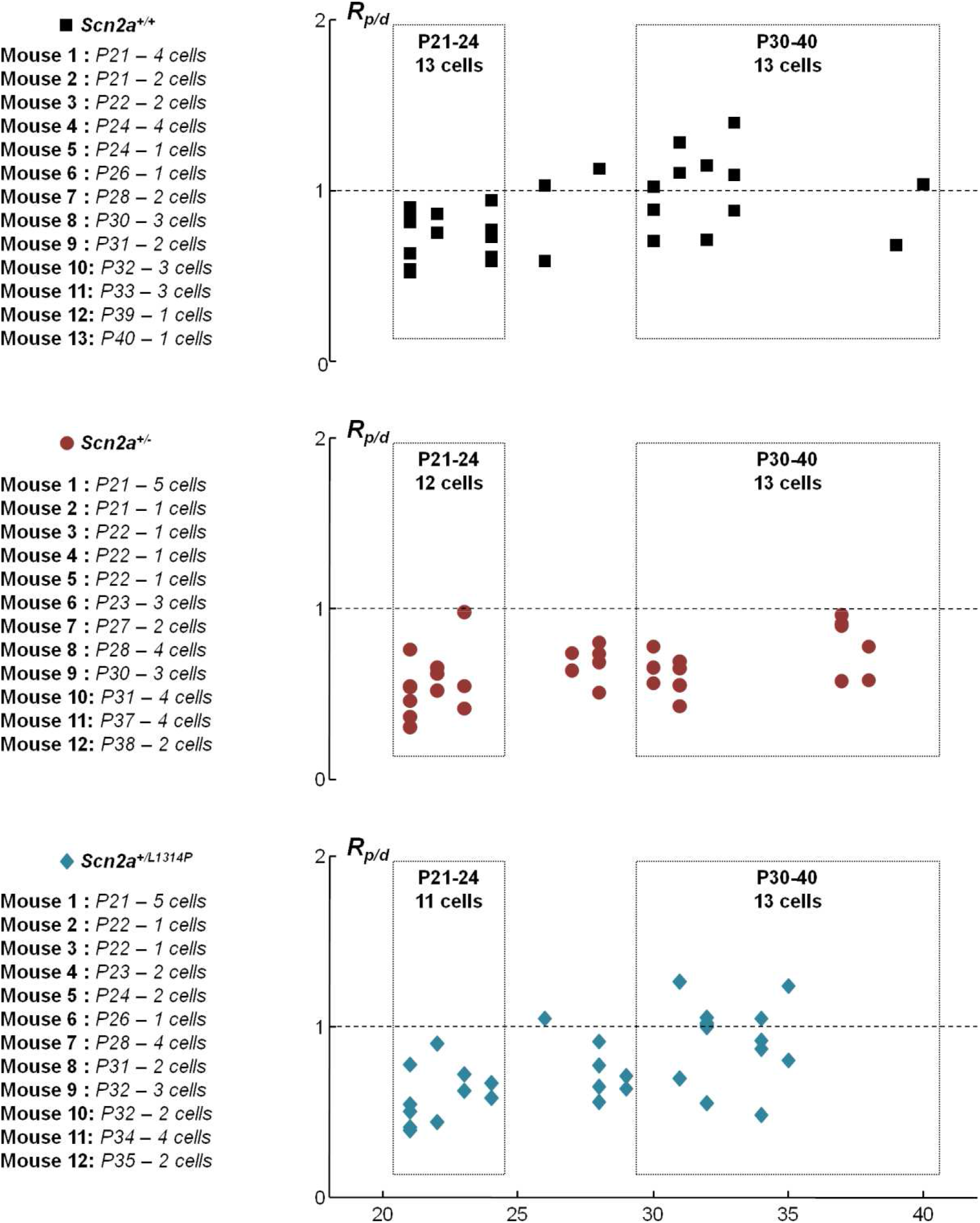
Scatter plot of R_p/d_ against age in Na^+^ influx measurements. Top, scatter plot of R_p/d_ against age of 29 *Scn2a^+/+^* cells from 13 female mice (table on the left). The groups “young” (P21-24) and “old” (P30-40) are outlined by dotted rectangles. *R_p/d_* has tendency of increasing with age (P=0.0076 and SP=0.757, ks-test between P21-24 and P30-40 groups). Middle, scatter plot of R_p/d_ against age of 31 *Scn2a^+/-^* cells from 12 female mice (table on the left) with no significant increase with age (SP=0.499 and P=0.0145, ks-test between P21-24 and P30-40 groups). Bottom, scatter plot of R_p/d_ against age of 29 *Scn2a^+/L1314P^* cells from 12 female mice (table on the left), with significant increase with age (P=0.0039 and SP=0.970, ks-test between P21-24 and P30-40 groups).

**Supplemental Figure 2.**
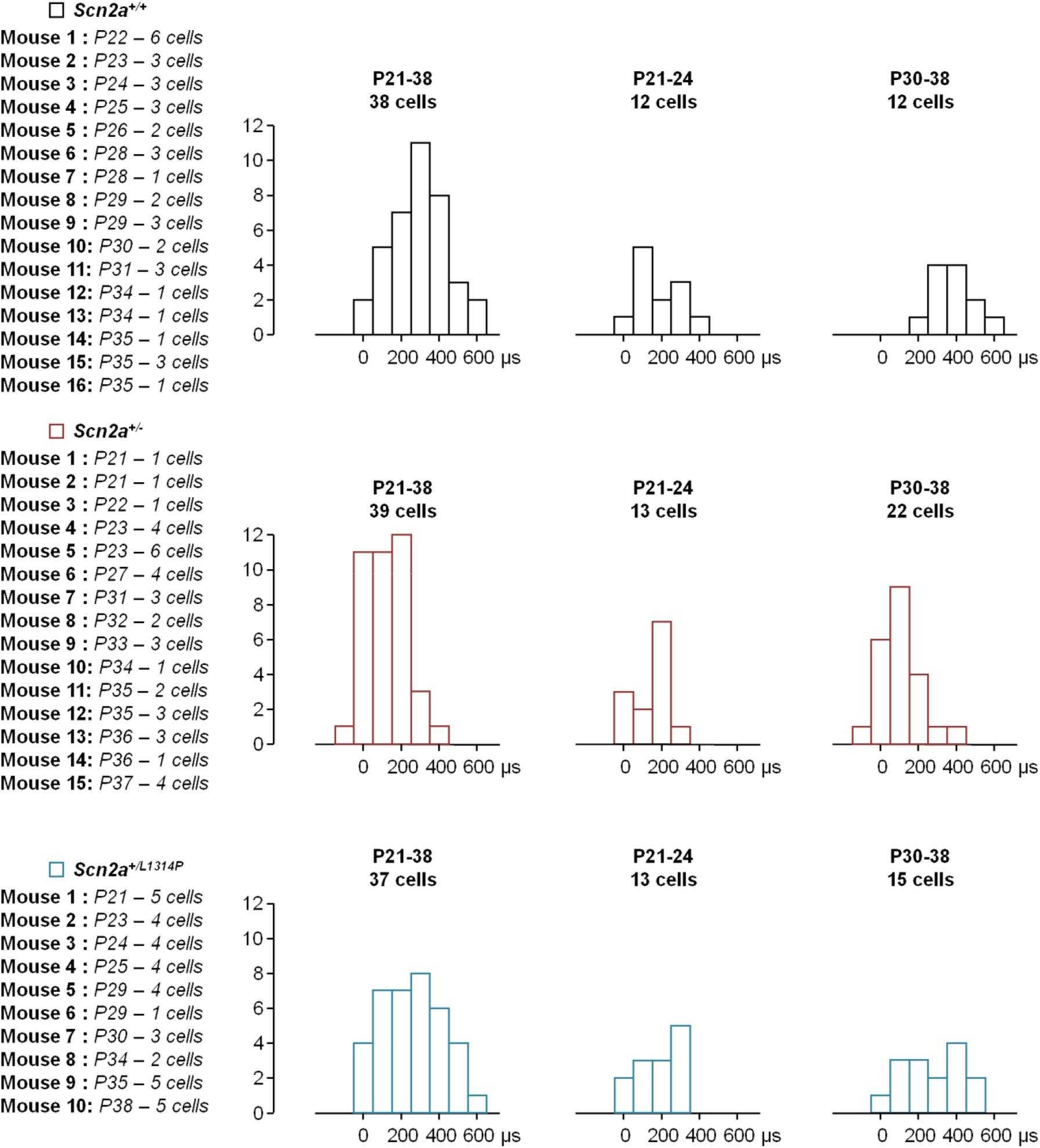
Histograms of delays between peaks of the Ca^2+^ current in proximal sites with respect to distal sites. Top, values from 38 *Scn2a^+/+^* cells from 16 female mice (table on the left) at P21-38, P21-24 (12 cells) and P30-40 (12 cells). Middle, values from 39 *Scn2a^+/-^* cells from 15 female mice (table on the left) at P21-38, P21-24 (13 cells) and P30-40 (22 cells). Bottom, values from 37 *Scn2a^+/L1314P^* cells from 10 female mice (table on the left) at P21-38, P21-24 (13 cells) and P30-40 (15 cells).

**Supplemental Figure 3.**
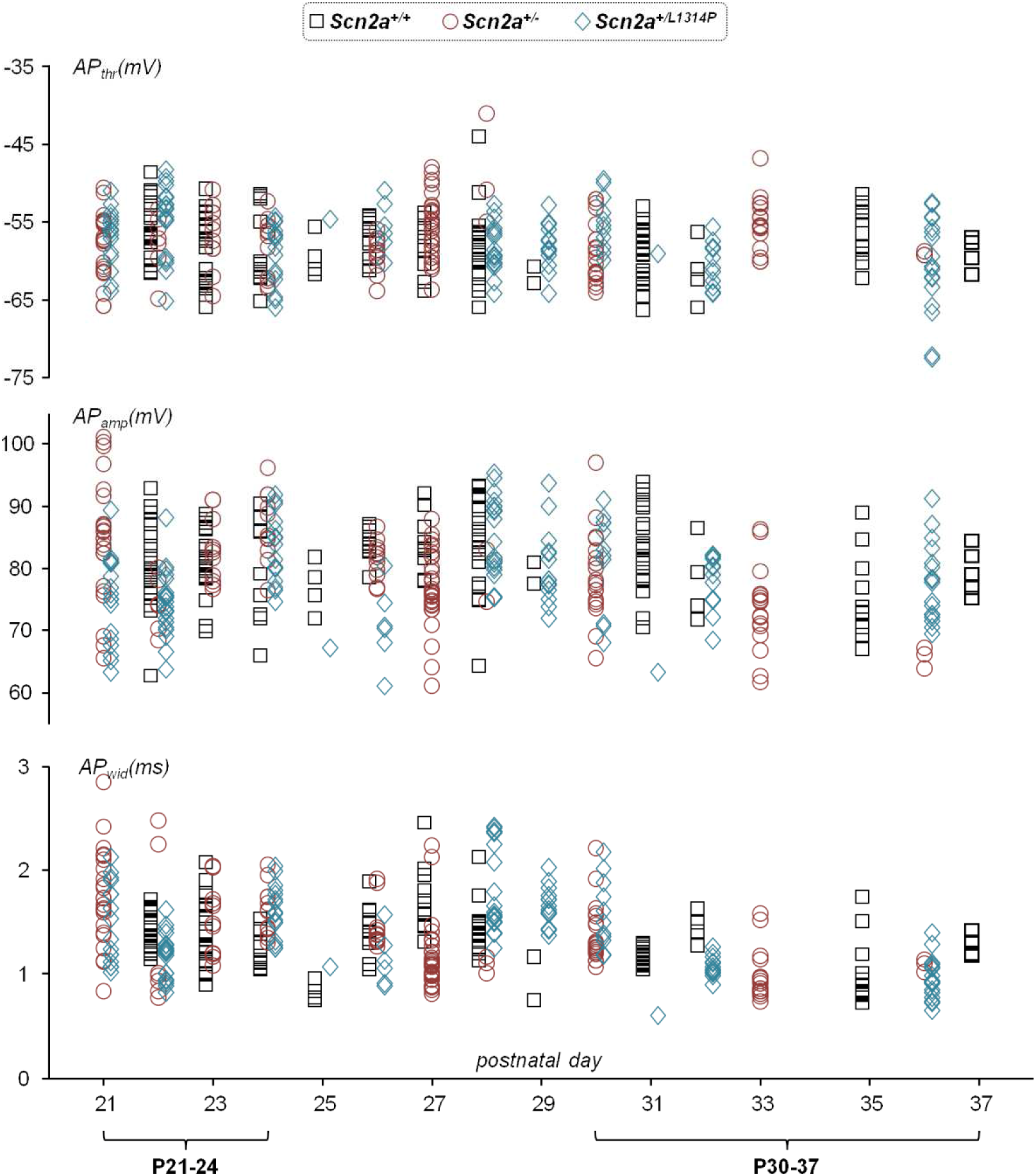
Scatter plots of analysed somatic potential parameters against age. Scatter plots of AP_thr_ (top), AP_amp_ (middle) and AP_wid_ (bottom) parameters in 142 *Scn2a^+/+^* cells, in 138 *Scn2a^+/-^* and in 136 *Scn2a^+/L1314P^* cells. The two ranges (P21-24 and P30-40) are indicated.

**Supplemental Figure 4.**
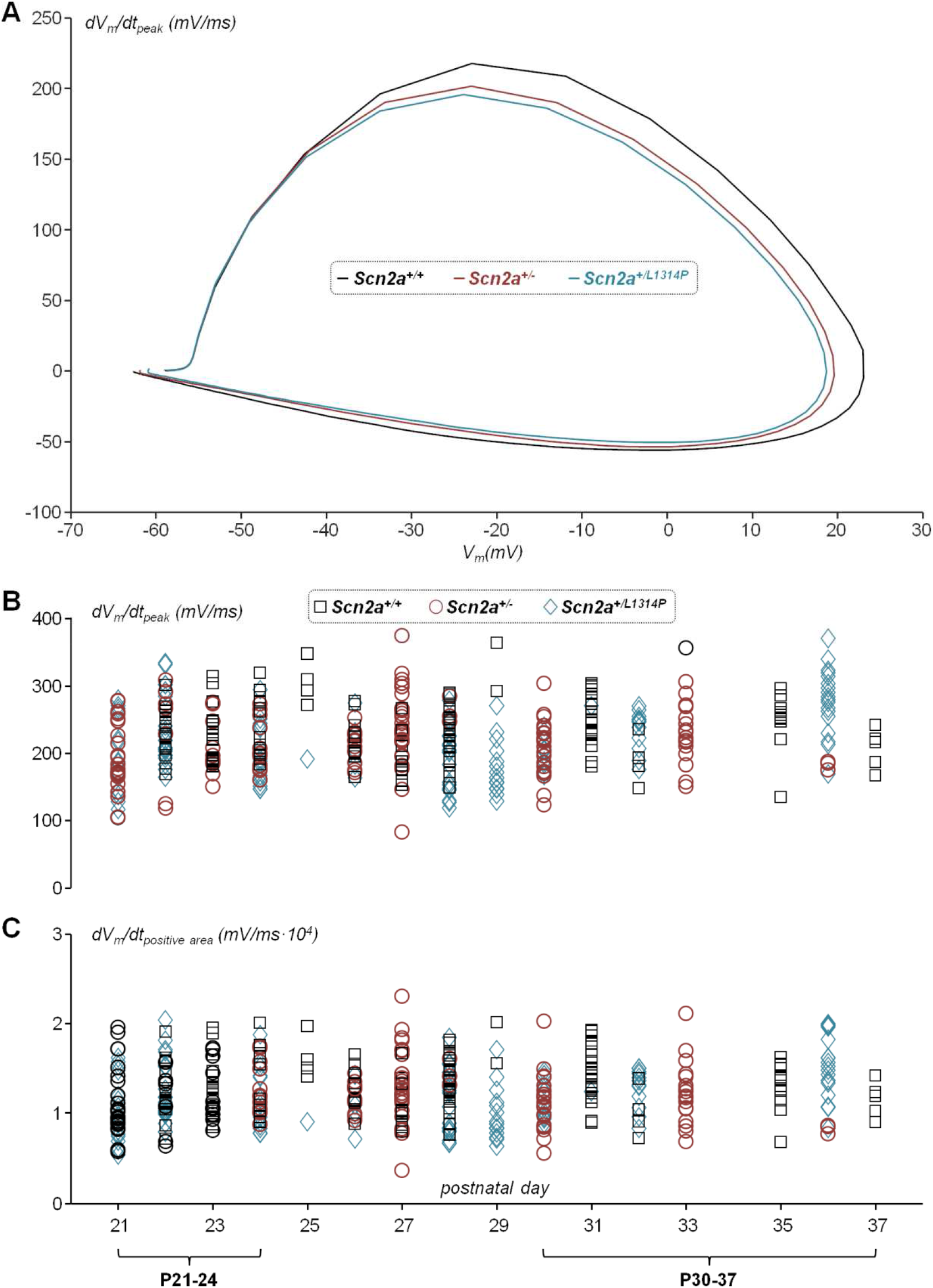
Analysis of the somatic V_m_ time derivative (*dV_m_/dt*). (A) *dV_m_/dt* versus V_m_ averaged over 142 *Scn2a^+/+^* cells (black trace), in 138 *Scn2a^+/-^* cells (brown trace) and in 136 *Scn2a^+/L1314P^* cells (light blue trace). (B) Scatter plot of *dV_m_/dt* peaks for the three strains. (C) Scatter plot of *dV_m_/dt* positive areas for the three strains.

**Supplemental Figure 5.**
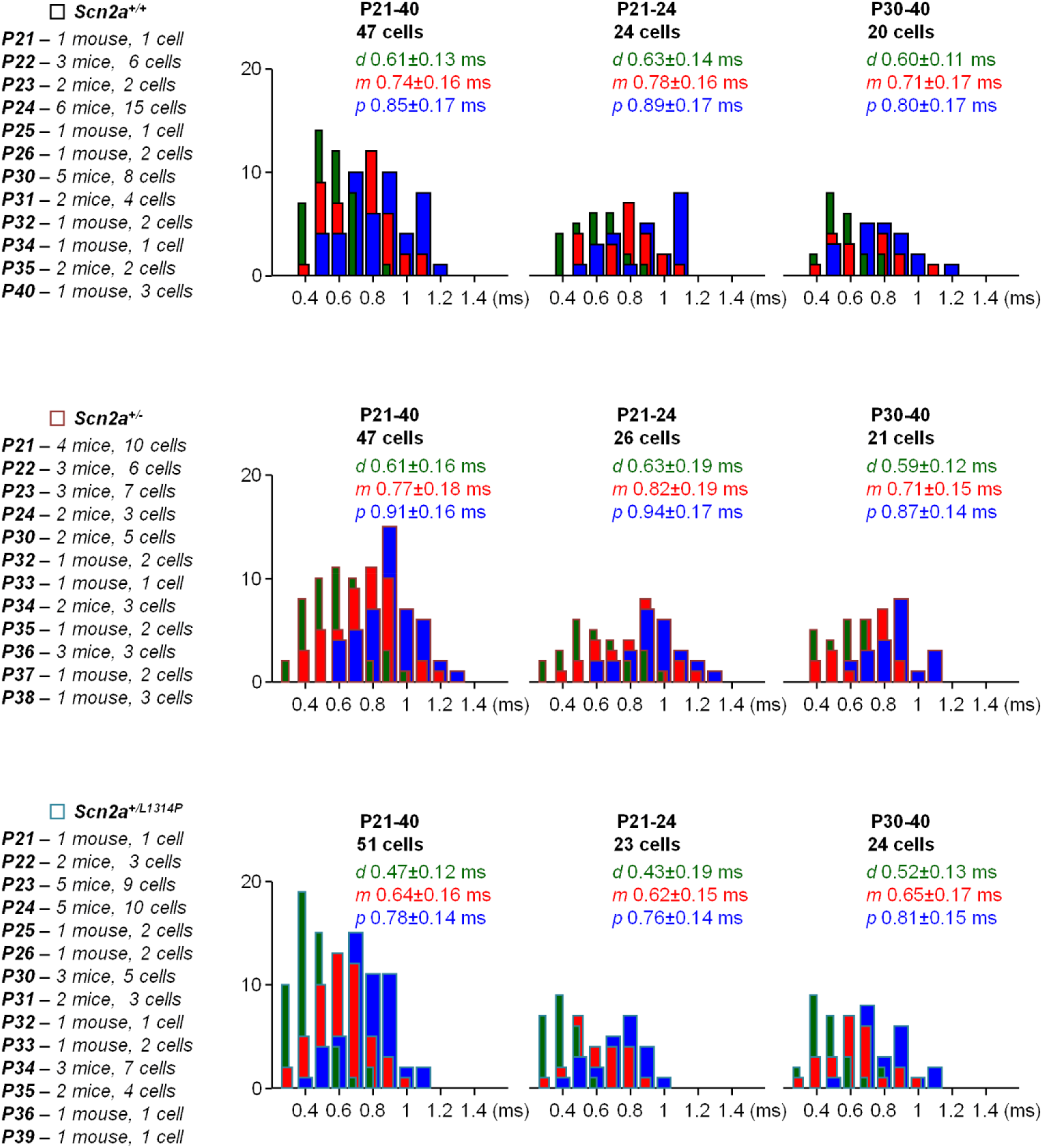
Histograms of AIS *APwid* at *d*, *m* and *p* sites. Top, *AP_wid_* histograms from 47 *Scn2a^+/+^* cells from 26 female mice (table on the left) at P21-40, P21-24 (24 cells) and P30-40 (20 cells). Middle, *AP_wid_* histograms from 47 *Scn2a^+/-^* cells from 24 female mice (table on the left) at P21-40, P21-24 (26 cells) and P30-40 (21 cells). Bottom, *AP_wid_* histograms from 51 *Scn2a^+/L1314P^* cells from 29 female mice (table on the left) at P21-40, P21-24 (23 cells) and P30-40 (24 cells). Mean ± SD of *AP_wid_* for each histogram is reported.

**Supplemental Figure 6.**
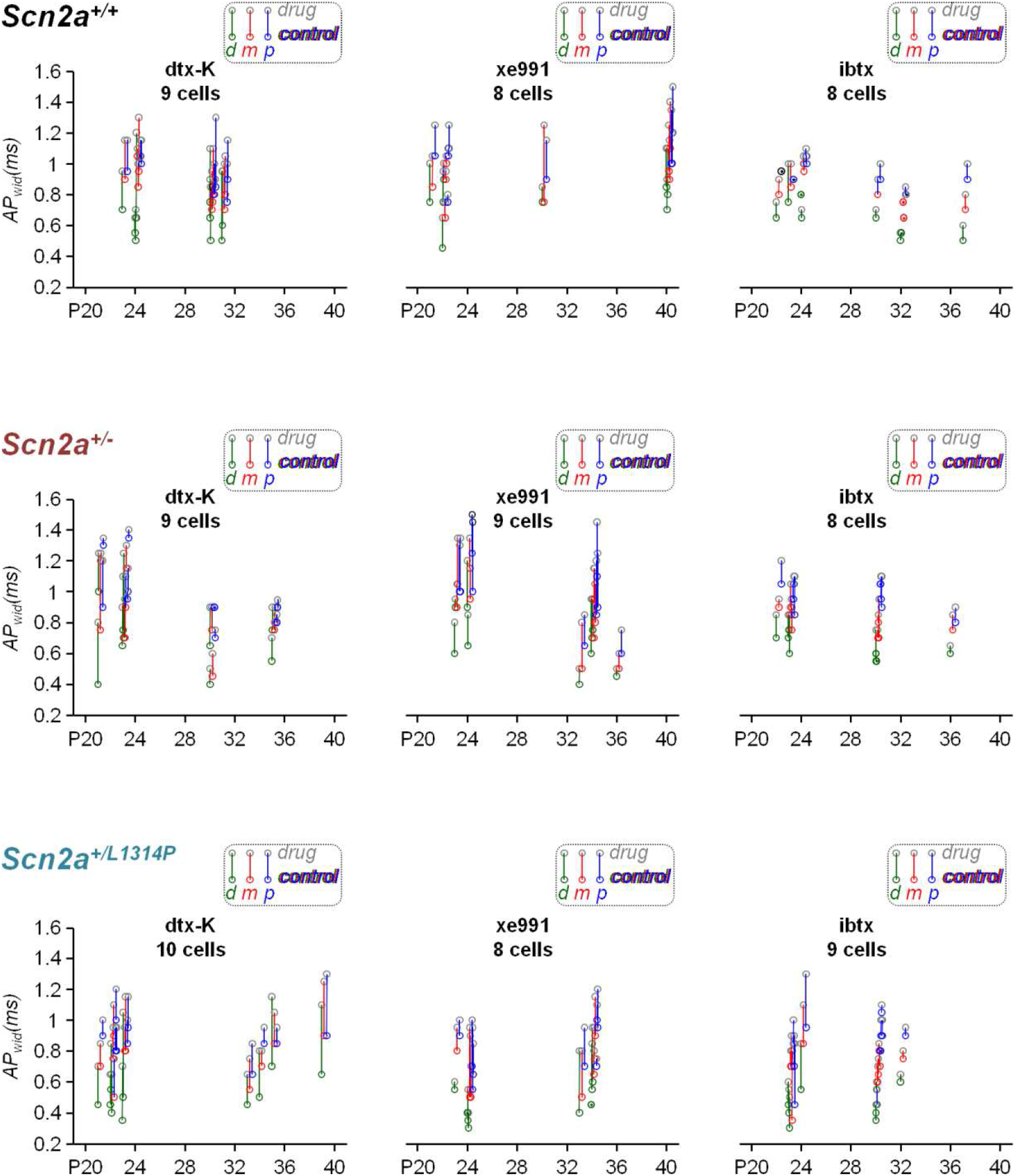
Scatter plots of AIS *APwid* fractional changes at *d*, *m* and *p* sites after delivery of dtx-K, xe991 or ibtx. Top-left, *AP_wid_* before and after delivery of 500 nM dtx-K against age in *Scn2a^+/+^* cells; top-centre, *AP_wid_* before and after delivery of 10 µM xe991 against age in *Scn2a^+/+^* cells; top-right, *AP_wid_*before and after delivery of 1 µM ibtx against age in *Scn2a^+/+^* cells. *AP_wid_* before and after delivery of 500 nM dtx-K against age in *Scn2a^+/-^* cells; middle-centre, *AP_wid_* before and after delivery of 10 µM xe991 against age in *Scn2a^+/-^* cells; middle-right, *AP_wid_*before and after delivery of 1 µM ibtx against age in *Scn2a^+/-^* cells. Bottom-left, *AP_wid_*before and after delivery of 500 nM dtx-K against age in *Scn2a^+/L1314P^* cells; bottom-centre, *AP_wid_* before and after delivery of 10 µM xe991 against age in *Scn2a^+/L1314P^* cells; bottom-right, *AP_wid_* before and after delivery of 1 µM ibtx against age in *Scn2a^+/L1314P^* cells. Coloured lines indicate *AP_wid_* increase from coloured circle to gray circle after drug delivery. Coloured dots indicate no *AP_wid_* change after drug delivery.

**Supplemental Figure 7.**
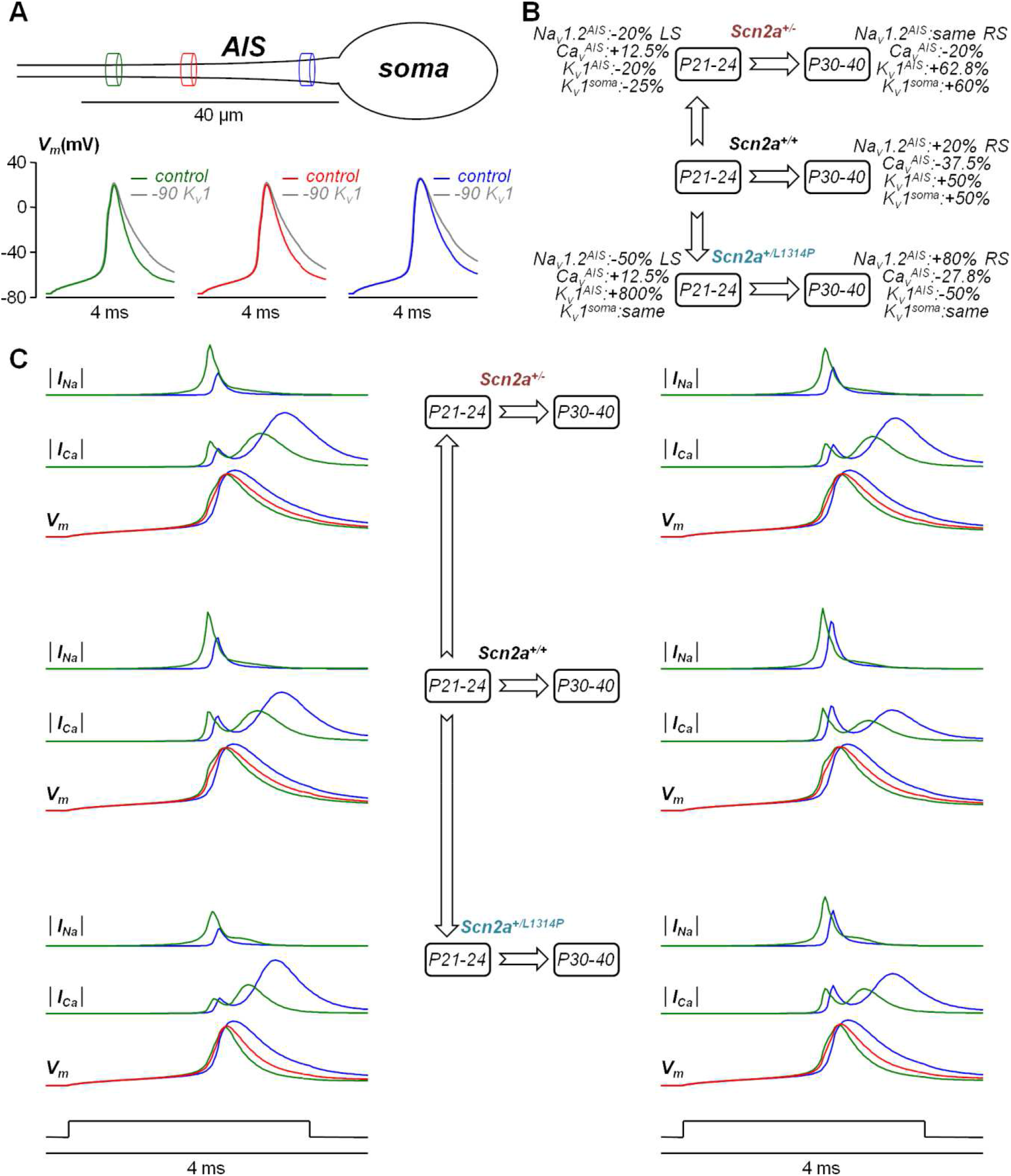
Complement of NEURON-simulations of “minimal” model mimicking juvenile AIS maturation in *Scn2a^+/+^, Scn2a^+/-^* and *Scn2a^+/L1314P^* L5 pyramidal neurons. (A) On the top, position of distal (*d*, green cylinder) medial (*m*, red cylinder) and proximal (*p*, blue cylinder) sites used to reproduce experimental results. On the bottom, V_m_ at *d, m*, and *p* sites associated with AP from the initial parameter corresponding to *Scn2a^+/+^* neuron (coloured traces) and with the same set but removing 90% K_v_1 from the AIS and the soma (grey traces). (B) Percentage changes of Na_v_1.2, K_v_1 and Ca_v_ densities (AIS or soma) from a set to the other pointed by the arrow; LS and RS mean, respectively, “left shift” and “right shift” of the distribution with respect to the scheme in panel A; simulations mimic P21-24 to P30-40 maturation in *Scn2a^+/+^, Scn2a^+/-^* and *Scn2a^+/L1314P^*L5 pyramidal neurons; (C) From the six simulations, absolute *I_Na_*(│*I_Na_* │) at *d* and *p* sites, absolute *I_Ca_*(│*I_Ca_* │) at *d* and *p* sites, and V_m_ at *d, m*, and *p* sites. These original traces were used to calculate those reported in Figure 6B-D.

